# Characterization of changes in the hemagglutinin that accompanied the emergence of H3N2/1968 pandemic influenza viruses

**DOI:** 10.1101/2021.04.19.439873

**Authors:** Johanna West, Juliane Röder, Tatyana Matrosovich, Jana Beicht, Jan Baumann, Nancy Mounogou Kouassi, Jennifer Doedt, Nicolai Bovin, Gianpiero Zamperin, Michele Gastaldelli, Annalisa Salviato, Francesco Bonfante, Sergei Kosakovsky Pond, Sander Herfst, Ron Fouchier, Jochen Wilhelm, Hans-Dieter Klenk, Mikhail Matrosovich

## Abstract

The hemagglutinin (HA) of A/H3N2 pandemic influenza viruses (IAVs) of 1968 differed from its inferred avian precursor by eight amino acid substitutions. To determine their phenotypic effects, we studied recombinant variants of A/Hong Kong/1/1968 virus containing either human-type or avian-type amino acids in the corresponding positions of HA. The precursor HA displayed receptor binding profile and high conformational stability typical for duck IAVs. Substitutions Q226L and G228S, in addition to their known effects on receptor specificity and replication, marginally decreased HA stability. Substitutions R62I, D63N, D81N and N193S reduced HA binding avidity. Substitutions R62I, D63N, D81N and A144G promoted virus replication in human airway epithelial cultures. Analysis of HA sequences revealed that substitutions D63N and D81N accompanied by the addition of N-glycans represent common markers of avian H3 HA adaptation to mammals. Our results advance understanding of genotypic and phenotypic changes in IAV HA required for avian-to-human adaptation and pandemic emergence.

## Introduction

Wild aquatic birds represent the major natural reservoir of IAVs, which occasionally transmit, adapt and circulate for prolonged periods of time in domestic birds and mammals (Olsen et al., 2006; Yoon et al., 2014). Because animal IAVs do not replicate efficiently in humans, zoonotic transmissions of IAVs are typically restricted to isolated cases of infection [for a recent review, see (Wang et al., 2020)]. If a zoonotic IAV against which people have no protective immunity acquires the ability to transmit efficiently in humans, it may initiate an influenza pandemic. Genetic and virological data available for the four last pandemic IAVs (H1N1/1918, H2N2/1957, H3N2/1968, and H1N1/2009) indicate that they all contained antigenically novel hemagglutinin (HA) gene segments derived from animal IAVs; the other gene segments originated from either animal or contemporary human IAVs [for reviews, see (Guan et al., 2010; Taubenberger and Kash, 2010)]. Thus, it is particularly important to understand which adaptive changes in the HA were required for the emergence of previous pandemic viruses from their animal precursors.

The HA mediates attachment of IAVs to sialic acid-containing glycan receptors on cells. Tropism, replication efficiency and pathogenicity of IAVs in different host species strongly depends on the optimal interplay between viral receptor-binding properties and spectra of sialoglycans expressed in target tissues of these species (reviewed by (Byrd-Leotis et al., 2017; de Graaf and Fouchier, 2014; Matrosovich et al., 2006b)). HAs of the previous pandemic IAVs differed from avian HAs by one or two amino acid substitutions in the conserved positions of the receptor-binding site (RBS). These substitutions were found to be essential for the switch of the HA receptor specificity from preferential binding to Neu5Acα2-3Gal-terminated glycans (avian-type receptors) to preferential binding to Neu5Acα2-6Gal-terminated glycans (human-type receptors). In the case of H2N2/1957 and H3N2/1968 IAVs, substitutions Q226L and G228S were responsible for this switch in receptor specificity. In the case of H1N1/1918 and H1N1/2009 IAVs, this role was played by substitutions E190D and G225D/E [for recent reviews see (Gamblin et al., 2020; Thompson and Paulson, 2020)]. It remains unexplored whether other substitutions in the HA of pandemic IAVs were required for adaptation to receptors in humans, for example, by adjusting HA interactions with sub-terminal oligosaccharide parts of the receptors and/or modulating binding avidity.

After endocytosis and acidification of endosomes, the HA of IAVs undergoes a low-pH-triggered conformational transition that mediates fusion between the viral and endosomal membranes. The conformational stability of the HA determines both the pH range of viral-endosomal fusion and stability of the virus in the environment. There is growing evidence that the pH optimum of fusion and stability of the HA differ between IAVs from different host species and that these differences may affect viral host range, pathogenicity, airborne transmission and pandemic potential [reviewed by (Russell et al., 2018)]. Human IAVs typically have a lower fusion pH optimum (from 5.0 to 5.4) than swine IAVs and zoonotic poultry IAVs of the H5 and H7 subtypes (pH from 5.6 to 6.2). The HAs of the H1N1/1918, H2N2/1957 and H3N2/1968 pandemic IAVs had a pH optimum of fusion typical for human viruses (5.1-5.4) (Baumann et al., 2016; Galloway et al., 2013). The earliest isolates of the H1N1/2009 had a less stable HA (pH optimum of fusion 5.4-5.5), but more stable variants were selected during a few months of virus circulation in humans (Cotter et al., 2014; Russier et al., 2016). The fusion pH and stability of the immediate HA precursors of the pandemic viruses were not studied, and it remains obscure whether alterations of these properties played a role in pandemic emergence.

We previously studied adaptive changes in the HA of the pandemic IAV A/Hong Kong/1/1968 (H3N2), which differed from the inferred avian ancestor HA by eight amino acid substitutions. Introduction of avian-virus-like amino acids at positions 226 and 228 of the HA altered cell tropism, reduced replication efficiency in cultures of human airway epithelial cells and abolished transmission of the virus in experimentally infected pigs (Matrosovich et al., 2007; Van Poucke et al., 2013). A combination of avian-type amino acid reversions at five other HA positions impeded replication in human airway cultures and markedly impaired transmissibility in pigs (Van Poucke et al., 2015). These results confirmed the critical role of substitutions Q226L and G228S in the avian-to-human transmission of the H3 HA and suggested that at least some of the other substitutions contributed to the emergence of the H3N2 pandemic virus.

In this study, we wished to further characterize changes in the HA that accompanied its avian-to-human adaptation during generation of the 1968 pandemic IAVs. We also wished to identify which substitutions, in addition to substitutions Q226L and G228S, played a role in the adaptation to humans. To address these questions, we prepared a panel of 18 recombinant variants of A/Hong Kong/1/1968 (H3N2) containing either human-type or avian-type amino acids at HA positions that separated the H3N2/1968 viruses from their inferred avian ancestor. We compared these IAVs for their membrane fusion activity and stability, receptor-binding properties, replication efficiency in MDCK cells and cultures of human airway epithelial cells. We also analyzed patterns of evolution of H3 HA codons in question in IAVs from different host species.

## 2. Materials and Methods

### 2.1. Cells and wild type IAVs

Cultivation of all non-infected and virus-infected cell cultures was performed at 37°C in 5% CO_2_. MDCK cells, human embryonic kidney 293T cells, and human bronchial adenocarcinoma Calu-3 cells were propagated using Dulbecco’s modified Eagle medium (DMEM; Gibco) supplemented with 10% fetal calf serum (FCS; Gibco), 100 IU/ml penicillin and 100 μg/ml streptomycin (pen-strep), and 2 mM glutamine. DMEM containing pen-strep, 2 mM glutamine and 0.1% bovine serum albumin (PAA Laboratories) (DMEM-BSA) was used for the viral infections.

Differentiated cultures of primary human tracheobronchial cells (HTBE) were prepared as described previously (Matrosovich et al., 2004). In brief, primary HTBE cells (Lonza) were expanded on plastic in BEGM growth medium (Lonza) and stored in aliquots in liquid nitrogen. Thawed passage-1 cells were grown on membrane supports (12-mm Transwell-Clear; pore size, 0.4 μm; Corning) in a 1:1 mixture of BEGM with DMEM. After 1 week, the medium was removed from the upper compartment and cells were maintained in BEGM/DMEM mixture under air-liquid interface (ALI) conditions. Fully differentiated 5- to 8-week-old cultures were used for the experiments.

A/Hong Kong/1/1968 (H3N2) was provided by Earl Brown, University of Ottawa, Ottawa, Ontario, Canada and grown in MDCK cells. A/Mallard/Alberta/279/1998 (H3N8) and A/Ruddy turnstone/Delaware/2378/1988 (H7N7) were provided by Robert Webster, St. Jude Children’s Research Hospital, Memphis, TN, USA. The avian viruses were grown in 11-days-old embryonated hen’s eggs.

### 2.2. Plasmids and recombinant IAVs

Reverse genetics plasmid pHW2000 and pHW2000 plasmids containing gene segments of A/Puerto Rico/8/1934 (H1N1) (PR8) were provided by Richard Webby and Robert Webster, St. Jude Children’s Research Hospital, Memphis, TN, USA. The eight pHW2000 plasmids containing gene segments of HK/68, modified HA plasmids R2 and R5 and corresponding recombinant viruses were prepared previously (Matrosovich et al., 2007; Van Poucke et al., 2013).

Mutations were introduced into the HA plasmid of A/Hong Kong/1/1968 using a site-directed mutagenesis kit (QuikChange; Stratagene). 2:6 recombinant IAVs containing wt and modified HA of A/Hong Kong/1/1968, NA of A/Hong Kong/1/1968 and the remaining six gene segments of PR8 were generated by reverse genetics (Hoffmann et al., 2000) as described before (Gerlach et al., 2017). These viruses and their designations are listed in the Fig. 1a. They were amplified in MDCK cells using DMEM-BSA medium containing 1 μg/ml of TPCK-treated trypsin (Sigma), clarified by low-speed centrifugation, and stored in aliquots at −80°C. The identities of the HA- and NA-encoding genes of all viruses were confirmed by sequencing.

**Fig. 1.**
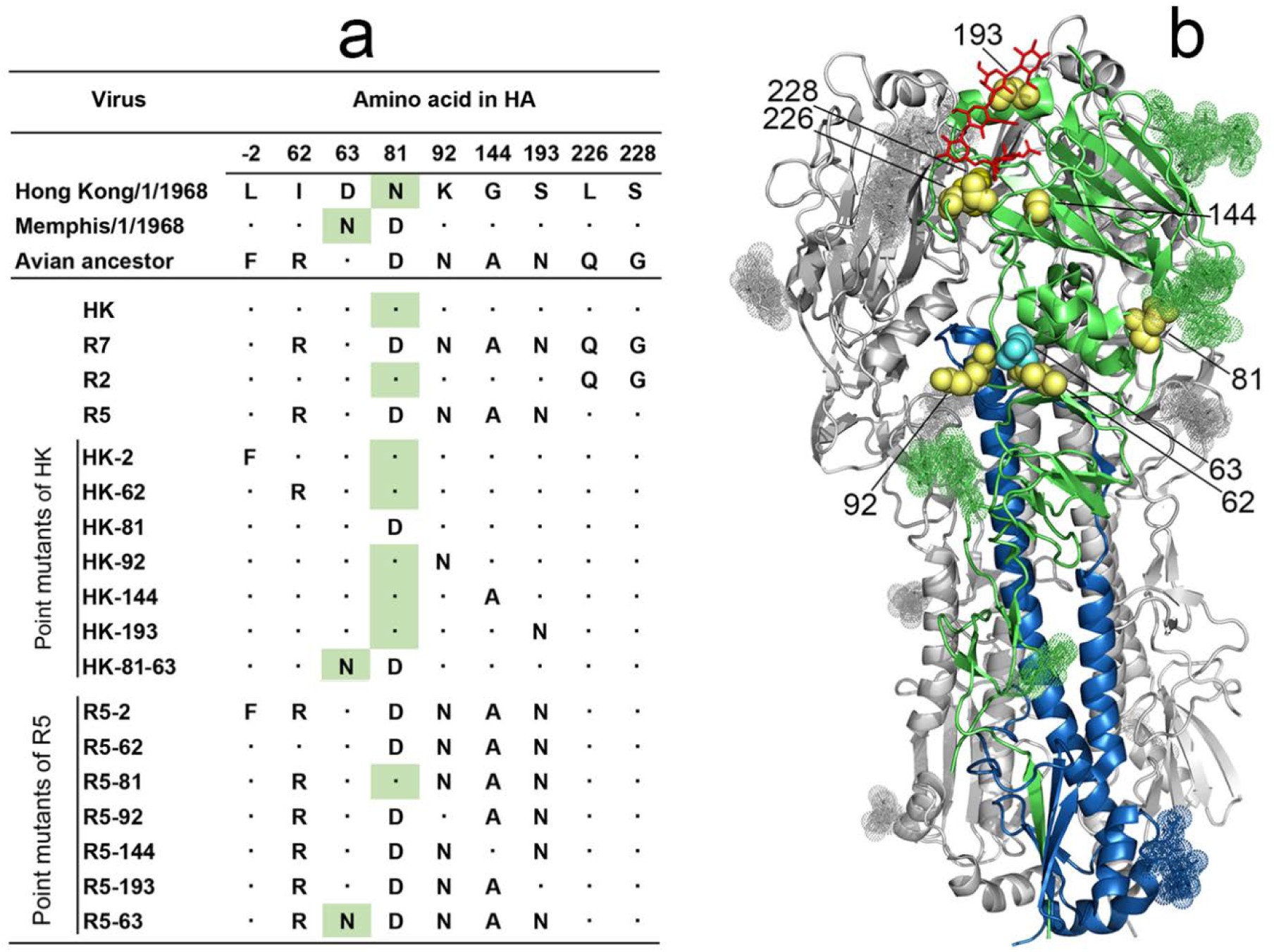
HA sequences and designations of 2:6 recombinant IAVs used in this study. (**a**) Amino acid differences between HAs of two 1968 pandemic virus lineages, their putative avian ancestor and 2:6 recombinant PR8-based viruses. Dots depict sequence identity with the HA of A/Hong Kong/1/1968. Numbering of amino acid positions starts from the N-terminus of the mature protein. Green background marks asparagine residues of glycosylation sites 63-65 and 81-83. (**b**) Location of amino acid substitutions shown as yellow space-filling models on the X-ray structure of the H3 HA complex with human receptor analogue LSTc (2YPG.pdb) (Lin, Xiong et al. 2012). Two HA monomers are colored gray, and the third monomer is colored green (HA1) and blue (HA2). LSTc is shown as red stick model, N-linked glycans are shown as dotted space-filling models. Cyan spheres show location of N_63_ present in the HA of A/Memphis/1/1968 lineage. The model was generated using PyMOL 2.0.6 (Schrödinger, LLC).

### 2.3. Virus titration and plaque size

Viruses were titrated in MDCK cells using single-cycle focus formation assay in 96-well plates (Matrosovich et al., 2007) and plaque formation assay under Avicel RC/CL overlay medium in 6-well plates (Matrosovich et al., 2006a). Infected cells were detected by immunostaining for viral nucleoprotein (NP). The viral concentrations were expressed in focus forming units (FFU) and plaque forming units (PFU) per ml, respectively. To determine the size of the plaques, plate wells containing from 5 to 50 plaques were scanned with a flat-bed scanner. The plaque diameters were measured with the Ruler Tool of Adobe Photoshop CS3 software version 10.0.1.

### 2.4. Low-pH-induced conformational transition of HA

Alteration of the HA sensitivity to protease digestion that accompany acid-induced conformational transition was determined using a solid-phase receptor binding assay as described previously (Matrosovich and Gambaryan, 2012; Van Poucke et al., 2015). In brief, viruses were adsorbed in the wells of fetuin-coated microtiter plates and incubated with buffers containing 25 mM MES, 150 mM NaCl, 0.9 mM CaCl_2_ and 0.5 mM MgCl_2_ (MES-NaCl). The pH of the buffers varied from 4.8 to 6.0 in 0.1 steps. After incubation with MES-NaCl buffers for 15 min at 37°C, the plates were washed with 25 mM phosphate buffered saline pH 7.2 (PBS) and incubated with 0.1 mg/ml of proteinase K in PBS for 1 h at 37°C. After washing with PBS containing 0.01% tween 80, binding of peroxidase-labelled fetuin (fet-HRP) was determined and expressed in percentages of binding to low-pH-exposed virus with respect to that of the virus exposed to pH 7. Binding-versus-pH curves were plotted, and pH values that corresponded to HA inactivation by 50% (pH_50_) were determined by linear interpolation.

### 2.5. Inactivation of HA by chaotropic agent and heat treatment

The effects of guanidinium hydrochloride (GnHCl) and elevated temperature on HA inactivation were quantified using the solid phase receptor binding assay described above. Viruses absorbed in the wells of fetuin-coated microtiter plates were either incubated for 1 h at 4°C with PBS containing variable concentrations of GnHCl or incubated with PBS for different time periods at 65°C. After washing, the binding of fet-HRP to GnHCl-treated and heat-treated viruses were determined and expressed in percentages with respect to the binding to control viruses incubated at 4°C with PBS. Concentration of GnHCl and incubation time at 65°C that reduced fet-HRP binding by 50% (IC_50_ and t_50_, respectively) were determined by linear interpolation.

### 2.6. Reduction of viral infectivity after 2-h incubation at 45 °C

Viral stocks were diluted in DMEM-BSA to a concentration of 4000 FFU per ml. Replicate 0.7-ml aliquots were incubated in closed Eppendorf tubes for 2 h either in a water bath at 45°C or on ice (control). All samples were next titrated using single-cycle focus formation assay in MDCK cells. Five technical replicates were used for the titration of each sample, and the results were averaged. The titers of heat-treated viruses were expressed as percentages of the corresponding control titers.

### 2.7. Low pH-induced polykaryon formation in virus-infected cells

pH-dependence of virus-induced cell-cell fusion was assayed as described (Reed et al., 2009) using MDCK cells instead of VERO cells. In brief, MDCK cultures in 96-well plates were inoculated with 1 FFU of the virus per cell in DMEM-BSA and incubated overnight. The medium was discarded, and the cultures were incubated for 15 min at 37°C with the DMEM-BSA containing 1 μg/ml of TPCK trypsin. The trypsin-containing medium was substituted by pre-warmed MES-NaCl pH-buffers (pH range from 5.3 to 7.0), incubated for 10 min at 37°C and washed once with PBS containing 0.9 mM CaCl_2_ and 0.5 mM MgCl_2_ (PBS+). After 3-h incubation with DMEM-BSA at 37°C, the cells were fixed with 70% ethanol, stained with Giemsa stain (Sigma) and analysed under the microscope. The highest pH at which more than 10 syncytia with more than 5 nuclei/syncytium were observed was taken as the pH threshold of polykaryon formation.

### 2.8. Infection inhibition by ammonium chloride

The assay determined virus dependence on endosomal acidification during infection as described previously (Baumann et al., 2016). In brief, MDCK cells in 96-well plates were inoculated with 200 FFU of the virus in 0.1 ml DMEM-BSA containing various concentrations of NH_4_Cl. The cells were incubated overnight, fixed and immuno-stained for viral NP. Concentrations of NH_4_Cl that reduced numbers of infected cells by 50% (IC_50_) were determined from dose-response curves by linear interpolation.

### 2.9. Infection inhibition by Vibrio cholerae sialidase

Binding avidity of the viruses for receptors on MDCK cells was compared using gradual desialylation of receptors with bacterial sialidase as described previously (Van Poucke et al., 2015). In brief, MDCK cells in 96-well plates were incubated with 0.05 ml per well of serial dilutions of sialidase in DMEM-BSA for 30 min at 37°C. Two hundred FFU of the viruses in 0.05 ml of DMEM-BSA were added per well without removing sialidase. No trypsin was added to the medium to avoid multicycle replication. The cultures were incubated overnight, fixed and immuno-stained for viral NP. Concentrations of sialidase that reduced numbers of infected cells by 50% (IC_50_) were determined from dose-response curves by linear interpolation.

### 2.10. Viral binding to sialoglycopolymers

Receptor-binding specificity of the viruses was characterized using soluble synthetic sialoglycopolymers (SGPs) (GlycoNZ, Auckland, New Zealand) (Tuzikov et al., 2021). The SGPs contained 20 mol% of sialyloligosaccharide moieties and 5 mol% of biotin attached to either the low-molecular-mass (20-kDa) or high-molecular-mass (1000-kDA) poly-N-(2-hydroxyethyl)acrylamide backbone.

The structures of the sialyloligosaccharide moieties and designations of SGPs are shown below.

|  |  |
| --- | --- |
| Neu5Acα2-3Galβ1-4GlcNAcβ | 3'SLN |
| Neu5Acα2-3Galβ1-4(6-Su)GlcNAcβ | 6-Su-3'SLN |
| Neu5Acα2-3Galβ1-4(Fucα1-3)GlcNAcβ | SLe<sup>x</sup> |
| Neu5Acα2-3Galβ1-4(Fucα1-3)(6-Su)GlcNAcβ | 6-Su-SLe<sup>x</sup> |
| Neu5Acα2-3Galβ1-3GlcNAcβ | SLe<sup>c</sup> |
| Neu5Acα2-3Galβ1-3GalNAcα | 3'STF |
| Neu5Acα2-6Galβ1-4GlcNAcβ | 6'SLN |

The binding of the viruses to SGPs was determined in a direct solid-phase binding assay as described previously (Matrosovich and Gambaryan, 2012). In brief, viruses adsorbed in the wells of fetuin-coated 96-well plates were allowed to interact with serially diluted SGPs followed by incubation with peroxidase–labelled streptavidin and tetramethylbenzidine (TMB) substrate solution. The association constants of virus complexes with SGPs (K_ass_) were determined from the slopes of A_450_/C versus A_450_ plots, where C is the concentration of the sialic acid in solution and A_450_ is the absorbance in the corresponding well.

### 2.11. Virus attachment and single-cycle infection in differentiated HTBE cultures

One day before the experiments, apical sides of the cultures were incubated with 0.15 ml of DMEM for 1 h at 37°C to collect secreted mucus. The mucus suspension was clarified by centrifugation at 6000x*g* for 5 min and stored at 4°C. Immediately before the experiments, the cultures were washed 10 times with PBS+.

To study virus attachment, the cultures were incubated with virus suspensions in DMEM-BSA for 1 h at 4°C. Control cultures were incubated with DMEM-BSA. The cultures were washed with PBS+ and fixed with 4% paraformaldehyde for 30 min at 4°C. Attached viruses were quantified by immunostaining of the apical sides of the cultures directly on Transwell-Clear supports. The cultures were blocked with 5% normal donkey serum (NDS, Dianova) at 4°C overnight, followed by sequential 1-h incubation at room temperature with in-house made rabbit polyclonal antibodies against HK and peroxidase-labelled donkey anti-rabbit antibodies (Dianova). Both antibodies were diluted in PBS buffer containing 10% normal horse serum (Dianova), 1% BSA, 1% NDS, 2% of the HTBE mucus suspension and 0.05% tween 80. After washing with 0.05% tween 80 in PBS, peroxidase activity was determined using TMB substrate. The mean substrate absorbency at 450 nm in the control cultures was subtracted from the absorbencies in virus-treated cultures.

To quantify concentration of physical virus particles in suspensions used in HTBE attachment experiments, non-specific virus binding to plastic was measured (Gambaryan et al., 1998a). Viral stocks were serially diluted in PBS and incubated in the wells of the immunoassay 96-well microplate (Greiner) overnight at 4°C (0.05 l/well). The wells were washed, fixed and immuno-stained with anti-HK antibodies and TMB substrate as described above for the HTBE experiments.

To study ability of the virus to enter into cell and initiate the first round of infection, replicate HTBE cultures were inoculated with 2×10^4^ FFU of the viruses in 0.2 ml of either DMEM-BSA or DMEM-BSA mixture with the mucus suspension collected the day before (3:1, vol/vol). The inoculums was removed 1 h post inoculation. The cultures were incubated for an additional 7 h at 37°C under ALI conditions, fixed, immuno-stained for viral NP, and infected cells were counted under an inverted microscope as described elsewhere (Gerlach et al., 2017).

### 2.12. Competitive replication in HTBE cultures

Competition between two viruses was studied by inoculating 5-6 replicate HTBE cultures with a mixture containing 4×10^3^ FFU of each virus in 0.2 ml DMEM. After 1-h incubation at 37°C, the inoculum was removed, and the cultures were incubated under ALI conditions. At 24, 48, 72, and 96 h post-inoculation, 0.3 ml DMEM was added to the apical sides of the cultures for 30 min. The apical medium was collected, stored at −80°C, and analysed together with the stored aliquot of the original inoculated virus mixture. Proportions of each HA genotype in the inoculated virus mixture and HTBE-harvests were determined by Sanger sequencing as described previously (Wendel et al., 2015).

Simultaneous competition between HK and 6 single-point HA mutants was studied as described above with the following modifications. HK and its 6 mutants were mixed in equivalent amounts based on infectious titers. Three different dilutions of this mixture were inoculated into HTBE cultures using 5 PFU of each virus per culture (low dose, L, 12 replicate cultures), 20 PFU per culture (medium dose, M, 12 cultures), and 320 PFU per culture (high dose, H, 6 cultures). The apical material was harvested once at 72 h post-inoculation and titrated for viral infectivity. From RNA extraction to variant calling, samples were processed as described previously (Wade et al., 2018), with the only exception of being sequenced for 300 bp paired-end. Proportions of each mutant HA genotype in the inoculated mixture and HTBE harvests were determined using the frequency of the single nucleotide polymorphism characterizing each mutant segment. In order to determine the effect of the inoculum titre on the proportion of each genotype in the harvest, we employed a generalized linear model relating proportions to inoculum titre (expressed by the variable “treatment” as L/M/H), specific genotype and the interaction of these two variables. Since proportions of the genotypes were clearly over dispersed, the model assumed a beta-binomial distribution in which the dispersion parameter σ was estimated as a function of “treatment” (Rigby and Stasinopoulos, 2005); for details of model construction see Supplementary Method). P values reflecting differences between the harvest and the inoculum were adjusted according to Dunnett’s method (Lenth et al., 2019).

### 2.13. Airborne transmission between ferrets

Respiratory droplet transmission experiments were performed as described previously (Munster et al., 2009). In brief, groups of two seronegative female adult ferrets were inoculated intranasally with 10^6^ TCID_50_ of each virus by applying 0.25 ml of virus suspension to each nostril. One day after inoculation, one naive ferret was placed opposite to each inoculated ferret in a transmission cage that prevented direct contact but allowed airflow from the inoculated to the naïve ferret. Nose and throat swabs were collected from inoculated and contact ferrets on days 1, 3, 5, and 7 post-inoculation and days 1, 3, 5, 7 and 9 post-exposure, respectively. Virus titers in swabs were determined by end-point titration in MDCK cells. Blood was collected from all ferrets on day 14 post exposure, and the presence of antibodies against the tested viruses was analysed by hemagglutination inhibition assay using standard procedures (WHO, 2002). All animals were humanely killed at the end of the in-vivo phase of the study.

### 2.14. HA sequences and phylogenetic analyses

Full-length nucleotide sequences of the H3 HAs were downloaded from the GISAID EpiFlu database (Shu and McCauley, 2017) accessed on March 11, 2020. Sequences were aligned using the MAFFT multiple alignment program implemented in the Unipro UGENE package (Okonechnikov et al., 2012), version 35. Sequences containing gaps and ambiguities, non-unique sequences and sequences of laboratory-derived IAVs were removed manually using Bio-Edit version 7.1.11 (Hall, 2004). Jalview version 2.11 (Waterhouse et al., 2009) was used to select representative sequences of swine IAVs with a redundancy threshold of 99%. The evolutionary history was inferred using IQ-TREE 2 with ModelFinder (Kalyaanamoorthy et al., 2017; Minh et al., 2020), the tree was plotted using MEGA7 (Kumar et al., 2016). Protein logos were generated using web-based application WebLogo (Crooks et al., 2004).

### 2.15. Selection pressure analyses

Three groups of host-specific HA sequences were analyzed, which included 1492 sequences of avian IAVs, 406 sequences of equine, canine, feline and seal IAVs and 803 sequences of human IAVs isolated in the years from 1968 to 1999. We partitioned the maximum likelihood tree into three groups of *internal* branches: human (801 branches), avian (1496 branches) and mammalian (394 branches).. Our analyses used dN/dS techniques [for a review, see (Pond et al., 2006)]. Because internal branches encompass at least one transmission event, we can assume that changes occurring along these branches have been “seen” by selection. Not including changes occurring along terminal branches we reduce the biasing effect of intra-host variation, which may be maladaptive on the population level (Pond et al., 2006), and tends to inflate dN/dS estimates (Kryazhimskiy and Plotkin, 2008). For each site in the HA alignment, we addressed the following four questions based on tests available in the HyPhy 2.5 package (Kosakovsky Pond et al., 2020).

1. What is the mean dN/dS at a site along the branches of interest? Does the site evolve subject to pervasive negative (dN/dS < 1) or positive diversifying (dN/dS > 1) selection? This test uses the Fixed Effects Likelihood (FEL) method (Kosakovsky Pond and Frost, 2005), and significance was established using a likelihood ratio test (LRT), at p ≤ 0.05. In addition, we inferred the number of synonymous and non-synonymous changes and the most likely character at each internal node of the tree using the SLAC method.
2. Does the site evolve subject episodic positive diversifying selection (dN/dS > 1 along some fraction of the tree)? This test used the Mixed Effects Model of Evolution (MEME) method (Murrell et al., 2012), and significance was established using LRT, at p ≤ 0.05.
3. Does the site evolve under different selective pressures between groups of branches (dN/dS differ between some or all of the four sets of branches)? This test used the Contrast Fixed Effects Likelihood (Contrast-FEL) method (Kosakovsky Pond et al., 2021), and significance was established using a collection of seven LRT (one for each pair of branch sets, and an omnibus test), with corrected p ≤ 0.01.
4. Is there evidence of directional evolution on “human” branches, where specific amino-acids are being selected for? This is based on an improved version of the Directional Evolution of Protein Sequences (DEPS) test (Kosakovsky Pond et al., 2008), and uses empirical Bayes Factors ≥ 100 to identify, which, if any residues at a given site are being selected for / against. This test is not based on dN/dS and is more suited to detect “sweeping” changes which involve only a few substitutions.

### 2.16. Statistics

Statistical tests were performed and using Graphpad Prism 8.4 and R 3.6.0 (www.r-project.org). Unless stated otherwise, figures show data from individual biological replicates. The bars or horizontal lines indicate the group means, the length of the error bars is one standard deviation. The details are explained in the table footnotes and figure legends. Student’s t test was used to compare two groups. Dunnett’s or Tukey’s multiple comparison tests were performed to compare more than two groups. More sophisticated statistical models were fitted by generalized linear models in R. If not stated otherwise, strictly positive variables were log-transformed before analysis, and percentage data were analyzed using quasi-binomial models with logit link. If the data consisted of experiments made on different days, day was included as a random intercept. Multiple tests of coefficients or contrasts within these models were done using simultaneous tests for general linear hypotheses, and P values were adjusted using the single step method (Hothorn et al., 2008) (comparable to Dunnett’s and Tukey’s procedure for multiple-to-one and all-pairwise comparisons). If mixed models were used to account for day-to-day variations between experiments, figures show data adjusted for day (that is, the variance attributed to day-to-day variation is removed from the data to better show how the values depend on the fixed factor). Details are explained in the methods sections of the respective experiments. Observed statistical significance is indicated in the figures as follows: *, P <0,05; **, P <0,01; ***, P <0,001.

## 3. Results

### 3.1. Preparation of recombinant variants of A/Hong Kong/1/1968-PR8 (H3N2) with substitutions in the HA

The 1968 pandemic IAV HA differed from the avian precursor by 8 amino acid substitutions (Bean et al., 1992; Van Poucke et al., 2015) (Fig. 1, supplementary Fig. S1). Seven substitutions were shared by all virus strains isolated in the first year of the pandemic. One of these substitutions, F(−2)L, was located in the cleavable signal peptide and six substitutions were located in the HA1 subunit of the mature HA protein, with G228S and Q226L in the RBS, A144G and N193S at the rim of the RBS and R62I and N92K in the vestigial esterase subdomain. The eighth substitution from D to N occurred at either position 63 or position 81 in the vestigial esterase subdomain of the HA and generated a new glycosylation site, N_63_-C_64_-T_65_ or N_81_-E_82_-T_83_, respectively. Either site was glycosylated with attached N-glycans detectable by X-ray analysis (see, for example, structures 4O58.pdb and 2YPG.pdb). Pandemic IAVs with these two HA variants differing solely by the location of new N-linked glycan co-circulated during 1968 and a few years afterwards. The A/Memphis/1/1968-like IAVs containing an N-glycan at position 63 (NG_63_) became extinct after 3 years of circulation; the A/Hong Kong/1/1968-like IAVs (NG_81_) continued to cause seasonal influenza outbreaks until 1976 and were substituted by a drift lineage that lost NG_81_ and gained NG_63_ (for the evolution of HA glycosylation sites 63 and 81 in human H3N2 viruses, see supplementary Fig. S2).

To study phenotypic effects of the substitutions, we generated a panel of 2:6 recombinant IAVs that contained HA and NA of A/Hong Kong/1/1968 (H3N2) and the remaining 6 gene segments of the laboratory strain PR8. The panel included the virus with wild type HA (HK) and its HA variants with either human-type or avian-type amino acids at corresponding HA positions (Fig. 1a). The R7 variant carried all seven avian-type substitutions in the mature HA, and thus mimicked the HA structure of the avian precursor of the 1968 pandemic IAVs. Two viruses were made to represent combined effects of substitutions at either positions 226 and 228 (variant R2) or at five other positions (variant R5). Single-point mutants of HK served to determine effects of reversions from human-type to avian-type amino acid at individual HA positions. The double mutant HK-81-63 represented the sequence of A/Memphis/1/1968 and was used to study the effect of the NG_63_. The point mutants of R5 were made to study effects of individual reversions from avian-type to human-type amino acids in the context of the avian HA with human-type L_226_ and S_228_. Finally, variants HK-2 and R5-2 were prepared to characterize the phenotype of the amino acid substitution in the signal peptide.

### 3.2. Effects of substitutions on HA conformational stability and membrane fusion activity

We first compared stability and fusion properties of HK, its avian precursor R7 and the intermediate variants R2 and R5 (Fig. 2). As these characteristics critically depend on the low-pH-triggered conformational transition of the HA, we determined the pH at which the viral HA changed its conformation by studying pH-induced alteration of HA sensitivity to protease digestion (Fig. 2a). R7 and R2 underwent conformational transition at a slightly lower pH than did HK and R5. In agreement with this finding, R7 and R2 initiated syncytia formation in MDCK cells at about 0.1 units of pH lower than did HK and R5 (Fig. 2b). To corroborate observed differences in viral fusion pH, we compared inhibition of viral infection in MDCK cells by ammonium chloride which counteracts endosomal acidification (Fig. 2c). R2 and R7 were more sensitive than HK and R5 to inhibition by NH_4_Cl, confirming that R2 and R7 require a lower pH in endosomes for fusion and cell entry.

**Fig.2.**
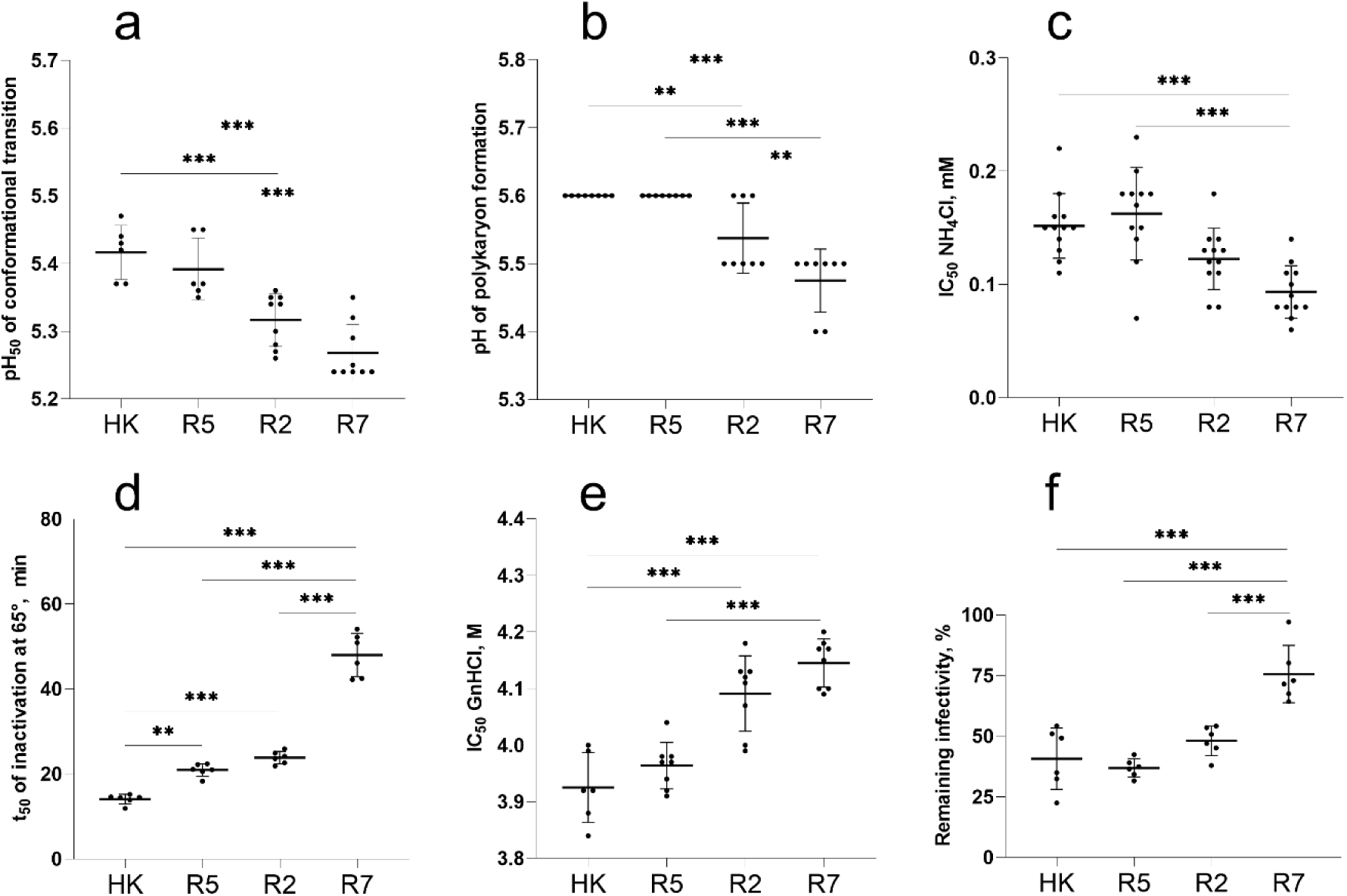
Conformational stability and membrane fusion properties of HK, R5, R2 and R7. (**a**) pH of acid-induced conformational transition of HA. Solid-phase adsorbed viruses were incubated in acidic buffers and treated with proteinase K. Viral binding of fet-HRP was assayed, and pH values that corresponded to 50% reduction of HA binding activity (pH_50_) were determined from binding-versus-pH curves. (**b**) pH threshold of polykaryon formation. Inoculated MDCK cells were cultured for 16 h, treated with trypsin and exposed to different pH buffers. After returning to neutral medium and incubation for 3 h, the cells were fixed, stained and analysed under the microscope. The data show highest pH values at which polykaryon formation was detected. (**c**) Inhibition of viral infection by ammonium chloride. MDCK cells were inoculated in the presence of various concentrations of NH_4_Cl, incubated overnight, fixed, and immunostained for NP. Concentrations of NH_4_Cl that reduced numbers of infected cells by 50% (IC_50_) were determined from dose-response curves. (**d**) HA stability at elevated temperature. Solid-phase adsorbed viruses were incubated in PBS at 65°C for different time periods and assayed for their binding to fet-HRP to determine incubation time required for 50% reduction of the binding activity (t_50_). (**e**) HA stability in chaotropic buffer. Solid-phase adsorbed viruses were incubated in buffers containing GnHCl for 60 min at 4°C washed with PBS and assayed for binding to fet-HRP. Data show concentrations of GnHCl that reduced viral binding activity by 50%. (**f**) Reduction of infectivity after incubation of the viruses for 2 h at 45°C determined by focus assay in MDCK cells. All panels show data points, mean values and SDs from 1 to 4 independent experiments performed with 2 to 7 replicates. P values for the differences between the viruses were determined with Tukey’s multiple comparison procedure.

Acid stability of the HA correlates, at least partially, with HA resistance to heat and denaturing agents. To test effects of the latter two factors of environmental stability we studied inactivation of the HA receptor-binding activity by the chaotropic agent guanidinium chloride (GnHCl) (Fig. 2e) and heat treatment (Fig. 2d) as well as the effect of heat treatment on viral infectivity (Fig. 2f). In general, the stability of the viruses in all three assays correlated with their acid stability. R7 was the most stable variant, HK was least stable, whereas R2 and R5 displayed intermediate stability.

We next studied the effects of non-226/228 single-point HA substitutions in HK and R5 using three assays (supplementary Fig. S3). None of the substitutions affected pH optimum of the HA conformational transition in the protease sensitivity assay. Four point mutants differed from the corresponding parental viruses by their sensitivity to ammonium chloride. Among them, mutants R5-63 and R5-81 containing N-linked glycan NG_63_ and NG_81_, respectively, were less sensitive than R5, whereas the mutant HK-81 lacking NG_81_ was more sensitive than HK. We concluded that N-glycan-containing variants entered the cells from less acidic endosomal compartment. This finding agrees with the study in which the presence of N-glycans in the globular head of H1N1 IAVs reduced receptor-binding avidity and facilitated HA-mediated fusion (Ohuchi et al., 2002). In the polykaryon formation assay, the avian-type amino acid in the signal peptide of the variants HK-2 and R5-2 correlated with a minor (ΔpH, +0.1) but reproducible elevation of their pH threshold of fusion (supplementary Fig. S3b). This effect, albeit small, was unexpected given that the signal peptide is not present in the mature HA and that HK-2 and R5-2 did not differ from their parents, HK and R5, in the conformational transition assay and in any other assays used (see below). Alteration of the signal peptide could potentially affect intracellular maturation, secretion and incorporation of the HA into virus particles, and recent bioinformatics analysis revealed that passaging of human IAVs in cell culture was occasionally accompanied by mutations in the signal peptide (Lee et al., 2019). These notions prompt further work on potential effects of the HA signal peptide on replication and interspecies adaptation of IAVs.

Collectively, this part of the study revealed that a combination of substitutions Q226L and G228S increased the pH of the conformational transition of the pandemic virus HA by about 0.15 pH units and as a consequence marginally decreased its environmental stability. The effects of other avian-to-human point substitutions and of their combination on conformational stability and membrane fusion activity were either smaller or below the detection limit of the assays.

### 3.3. Receptor-binding profile of the avian precursor of the 1968 pandemic viruses and effect of amino acid substitutions on the HA preference for the type of Neu5Ac-Gal linkage

To characterize receptor-binding properties of the viruses we determined their binding to soluble synthetic SGPs carrying multiple copies of sialyloligosaccharide moieties attached to a hydrophilic polymeric carrier. The high molecular mass (1 MDa) SGPs contained about 50 times more copies of the sialoligand per macromolecule and bound to IAVs with much higher avidity than structurally identical 20-kDa SGPs. As a result, utilization of the 1-MDa SGPs was instrumental for comparison of IAVs with large differences in binding avidity, such as avian and human IAVs, whereas the 20-kDa SGPs were more useful than 1-MDa SGPs for characterization of IAVs with minor differences in the binding avidity (Matrosovich and Gambaryan, 2012; Tuzikov et al., 2021).

Although most avian IAVs use Neu5Acα2-3Gal-terminated glycans as their cellular receptors, viruses adapted to species of the orders *Anseriformes*, *Charadriiformes and Galliformes* typically differ by their ability to recognize sub-terminal parts of the receptor glycans [(Gambaryan et al., 2018) and references therein]. We assumed that analysis of the fine receptor binding specificity of R7 may predict which avian species perpetuated the precursor of the 1968 pandemic virus. To this end, we determined binding of R7 and R2 to a panel of Neu5Acα2-3Gal-containing 20-kDa SGPs. Two representative wild-type viruses, A/mallard/Alberta/279/1998 (H3N8) (mal-H3N8) and A/ruddy turnstone/ Delaware/2378/1988 (H7N7) (rt-H7N7), were used for a comparison (Fig. 3a). R7 shared the binding profile with mal-H3N8 which represented typical receptor-binding properties of IAVs in ducks. Similar to other duck viruses (Gambaryan et al., 2018), R7 and mal-H3N8 bound poorly to fucosylated receptors SLe^x^ and 6-Su-SLe^x^ and bound more efficiently to SLe^c^ and 3’STF than to 3’SLN. A second control virus, rt-H7N7, displayed receptor-binding characteristics which are often shared by IAVs of *Charadriiformes and Galliformes* (Gambaryan et al., 2018; Gambaryan et al., 2012). Namely, this virus strongly bound to fucosylated receptors SLe^x^ and 6-Su-SLe^x^, bound to 3’SLN better than to SLe^c^, and bound particularly strongly to sulfated receptors 6-Su-3’SLN and 6-Su-SLe^x^. R7 did not display any of these features, thus showing no signs of adaptation of the R7 HA to gulls, shorebirds or gallinaceous poultry. The binding profile of the variant R2 was similar to the profiles of R7 and mal-H3N8 and only differed from these viruses by marginally elevated avidity for 3’SLN and 3’STF. Thus, five human-type amino acid substitutions separating R2 from R7 had only minor effect on HA binding to Neu5Acα2-3Gal-terminated SGPs and did not alter a typical duck-virus-like binding specificity of the HA.

**Fig. 3.**
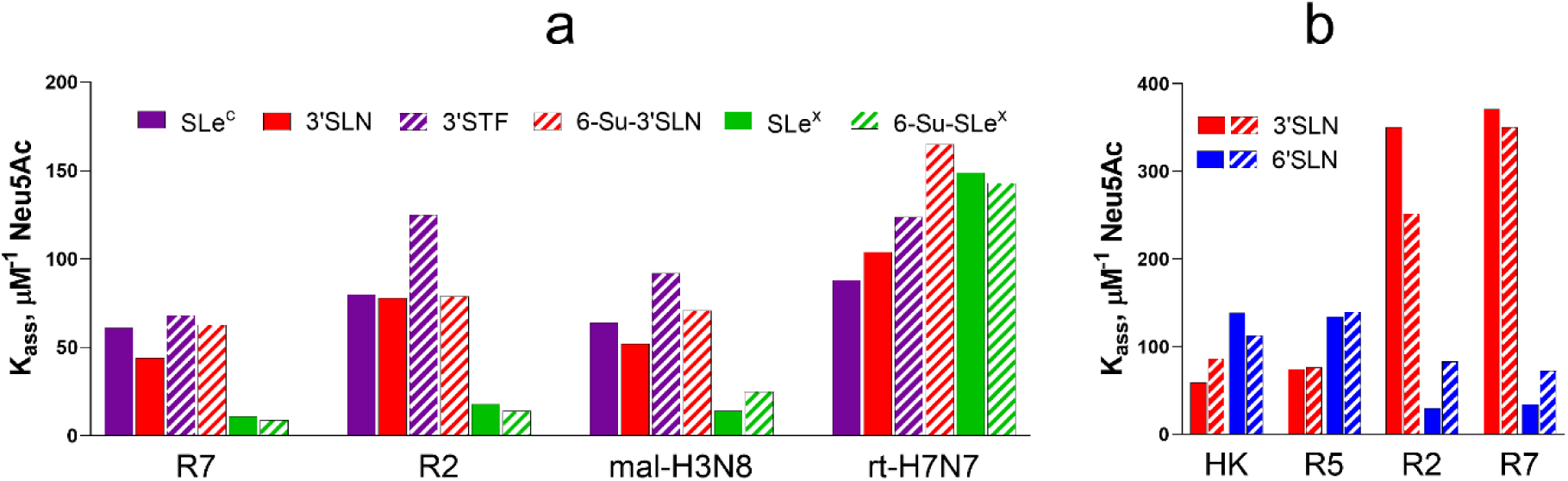
Binding of IAVs to biotinylated SGPs. The solid-phase adsorbed viruses were allowed to bind biotinylated SGPs from solution, and association constants of virus complexes with SGPs (K_ass_) were determined as described in Materials and Methods. Higher values of K_ass_ reflect stronger binding. (**a)** Binding to six low molecular mass Neu5Acα2-3Gal-containing SGPs differing by structure of penultimate sugar moieties. Wild type IAVs A/mallard/Alberta/279/1998 (H3N8) and A/ruddy turnstone/ Delaware/2378/1988 (H7N7) were tested in parallel with R7 and R2. Two to 4 experiments were performed on different days with similar results. Figure shows data from a representative experiment with one replicate for each virus-SGP. (**b)** Binding to high molecular mass SGPs 3’SLN and 6’SLN. Filled and hatched bars show mean values of experiments performed on two different days with two replicates per each virus-SGP.

Substitutions L226Q and G228S in the HAs of pandemic H3N2/1968 viruses switched the viral recognition of the type of Neu5Ac-Gal linkage (Matrosovich et al., 2006b; Thompson and Paulson, 2020). To determine whether and to what extent the other five substitutions in the mature HA contributed to this switch, we compared binding of HK, R5, R2 and R7 to 6’SLN and 3’SLN. High molecular mass SGPs were used to ensure measurable binding of each virus to both SGPs (Fig. 3b). As expected, comparison of R7 with R5 and comparison of R2 with HK showed that a combination of substitutions Q226L and G228S strongly reduced HA binding to 3’SLN and increased HA binding to 6’SLN; the magnitude of the second effect was noticeably smaller. No significant differences in the viral binding profiles were observed in pairs R7/R2 and HK/R5. These results indicated that a combination of substitutions at positions 62, 81, 92, 193 and 144 of the pandemic virus HA had much lower (if any) effect than substitutions Q226L/G228S on virus recognition of the Neu5Ac-Gal linkage type.

### 3.4. Effects of non-226/228 substitutions on binding avidity of the HA

As limited experiments with 1-MDa SGPs (Fig. 3b) did not reveal significant differences in their binding to HK and R5, we employed more sensitive receptor-binding assays to assess effects of amino acid substitutions separating these viruses. Fig. 4a shows data on binding of HK, R5 and their point mutants to low molecular mass 6’SLN. HK bound to 6’SLN about 2-fold weaker than R5 indicating that a combination of 5 human-type amino acids decreased binding avidity. Three single-point HK mutants, HK-62, HK-81 and HK-193 displayed elevated binding avidity. Three other HK mutants, HK-92, HK-144 and HK-2, did not differ from the parental HK. Remarkably, the double mutant HK-81-63 bound to 6’SLN significantly weaker than HK-81 and showed binding avidity that was comparable to that of HK. Thus, whereas substitution N81D and loss of NG_81_ increased HA binding to 6’SLN, the substitution D63N and attachment of NG_63_ fully compensated for this effect. The effects of human-type substitutions in R5 HA on virus binding to 6’SLN inversely correlated with the effects of corresponding avian-type substitutions in HK HA (Fig. 4a). Namely, substitutions at positions 62, 81, 193 and 63 in R5 decreased binding avidity, whereas substitutions at positions −2, 92 and 144 had no significant effect. This correlation indicated that the effects of individual substitutions on HA binding to 6’SLN do not depend on the identity of the other 4 amino acids studied (that is, avian-type amino acids in R5 and human-type amino acids in HK). We concluded that these amino acids are not involved in substantial epistatic interactions.

**Fig. 4.**
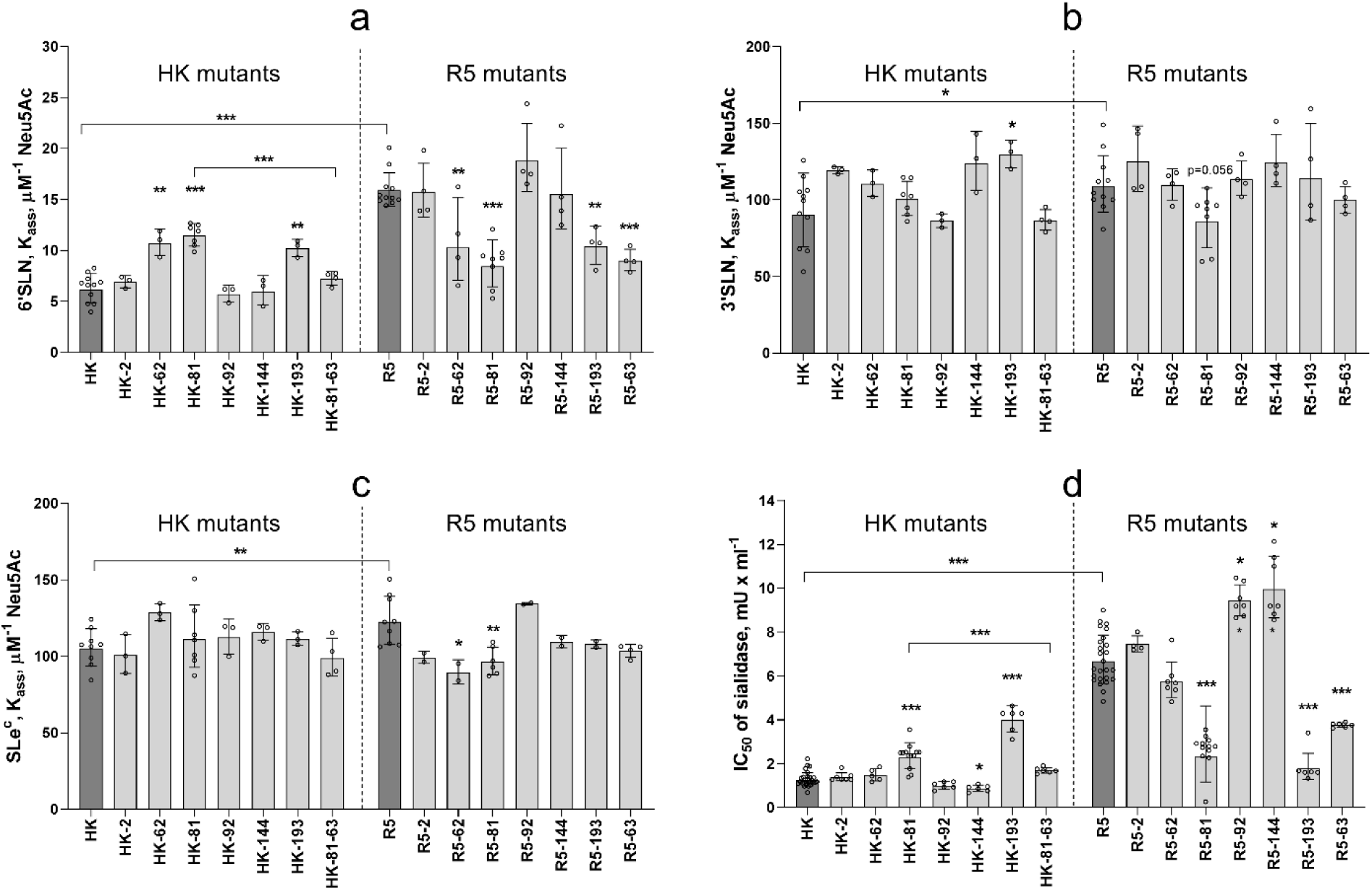
Receptor-binding properties of HA point mutants of HK and R5. (**a-c**) Association constants of viral complexes with biotinylated SGPs 6’SLN (20 kDa), 3’SLN and SLe^c^ (both 1 MDa) were determined as described in Materials and Methods. Data represent combined results from 4 to 11 experiments performed on different days with 1 replicate for each virus-SGP pair per experiment. (**d**) Inhibition of viral cell entry by *Vibrio cholerae* sialidase. MDCK cells were incubated with solutions of gradually diluted sialidase for 30 min, inoculated with 200 FFU of the viruses without removing sialidase, fixed after one cycle of replication and immunostained for viral NP. The figure shows concentrations of sialidase that reduced numbers of infected cells by 50% (IC_50_). From 2 to 9 experiments were performed on different days using 3 to 4 replicates per virus. All panels show the individual values adjusted for day as described in section 2.16 of Materials and Methods with geometric mean (bars) and SDs. Vertical dotted line separates point mutants of HK and point mutants of R5. Asterisks depict P values for the differences between single-point mutants and the corresponding parental virus, either HK or R5 (dark gray bars). Asterisks over horizontal lines depict differences between HK and R5 and between HK-81 and HK-81-63.

Because of the relatively weak binding of HK, R5 and point mutants to Neu5Acα2-3Gal-containing receptors, we could not reliably quantify binding of these IAVs to corresponding 20-kDa SGPs. Using more sensitive but less discriminative 1-MDa SGPs, we found that R5 bound stronger than HK to both 3’SLN and SLe^c^ and that most single-point mutants did not significantly differ in this respect from the parental IAVs (Fig. 4b,c). These results suggested that single-point substitutions had a weaker effect than their combination on HA binding to 3’SLN and SLe^c^.

To further characterize receptor-binding properties of the HA mutants, we studied inhibition of viral single-cycle infection in MDCK cells in the presence of *Vibrio cholerae* sialidase which reduced levels of sialic acid receptors on the cell surface (Fig. 4d). A higher value of 50% inhibitory concentration of sialidase (IC_50_) suggested that the virus can infect cells expressing lower amounts of receptor moieties; this effect was interpreted as an indication of a higher binding avidity. The avidity of the viruses for receptors on MDCK cells correlated to a large extent with viral binding to 6’SLN (compare Figs. 4a and 4d). Thus, in both assays, i) R5, HK-81 and HK-193 bound to the cells stronger than HK, ii) HK-81-63 bound weaker than HK-81, iii) R5-81, R5-193 and R5-63 bound weaker than R5, iv) substitution at position −2 of the signal peptide affected binding of neither R5, nor HK. In contrast with a significant effect of substitutions atn position 62 on binding to 6’SLN (Fig. 4a), these substitutions showed no apparent effect on binding to MDCK cells. As another distinction from the 6’SLN binding data, R5-92 and R5-144 bound to MDCK cells somewhat stronger than R5 and HK-144 bound weaker than HK.

The following conclusions could be made from the binding data. A combination of human-type substitutions separating HK from R5 reduced HA binding to both Neu5Acα2-6Gal-and Neu5Acα2-3Gal-containing receptors. Three of these substitutions, namely, R62I, N193S and either D81N or D63N, were primarily responsible for the reduction of binding avidity. The other three substitutions either had a weak binding-enhancing effect (N92K and A144G) or no effect [F(−2)L]).

The observed reduction of the avidity for human-type receptors during HA evolution from its avian precursor was unexpected. Since this result was obtained in experiments with MDCK-grown IAVs and with non-natural receptor analogues, we performed additional experiments using more natural experimental models. As shown previously, the relatively large N-glycans attached to the HA in MDCK cells could alter viral receptor-binding properties as compared to the same virus grown in another cell system (Gambaryan et al., 1998a; Inkster et al., 1993). To address this possibility, we re-grew a representative group of viruses in the human cell line Calu-3 and in differentiated cultures of primary human tracheal-bronchial epithelial cells (HTBE cultures). The Calu-3-grown HK, R5 and single-point mutants of HK displayed the same patterns of binding to 6’SLN and sialidase-treated cells (supplementary Fig. S4a,b) as did their MDCK-grown counterparts (Fig. 4a,d). The HTBE-grown HK, R5 and two glycosylation mutants R5-81 and R5-63 also showed the same relative binding avidity (supplementary Fig. S4c) as did corresponding MDCK-grown variants (Fig. 4d). These results indicated that MDCK-grown IAVs correctly represented receptor-binding phenotypes of the viruses during their replication (and glycosylation) in human epithelial cells.

To test whether differences in binding avidity of R5 and HK for SGPs and MDCK cells correlate with viral binding to biologically relevant receptors in humans, we studied attachment of R5 and HK to the apical surface of HTBE cultures which closely mimic structure and functions of human target cells in vivo (Davis et al., 2015; Matrosovich et al., 2004). R5 attached to HTBE cells more efficiently than HK (Fig. 5) in agreement with relative binding efficiency of these viruses to soluble 6’SLN and to MDCK cells. These results confirmed that non-226/228 human-type amino acid substitutions in the precursor avian HA reduced efficiency of virus binding to receptors on airway epithelial cells in humans.

**Fig. 5.**
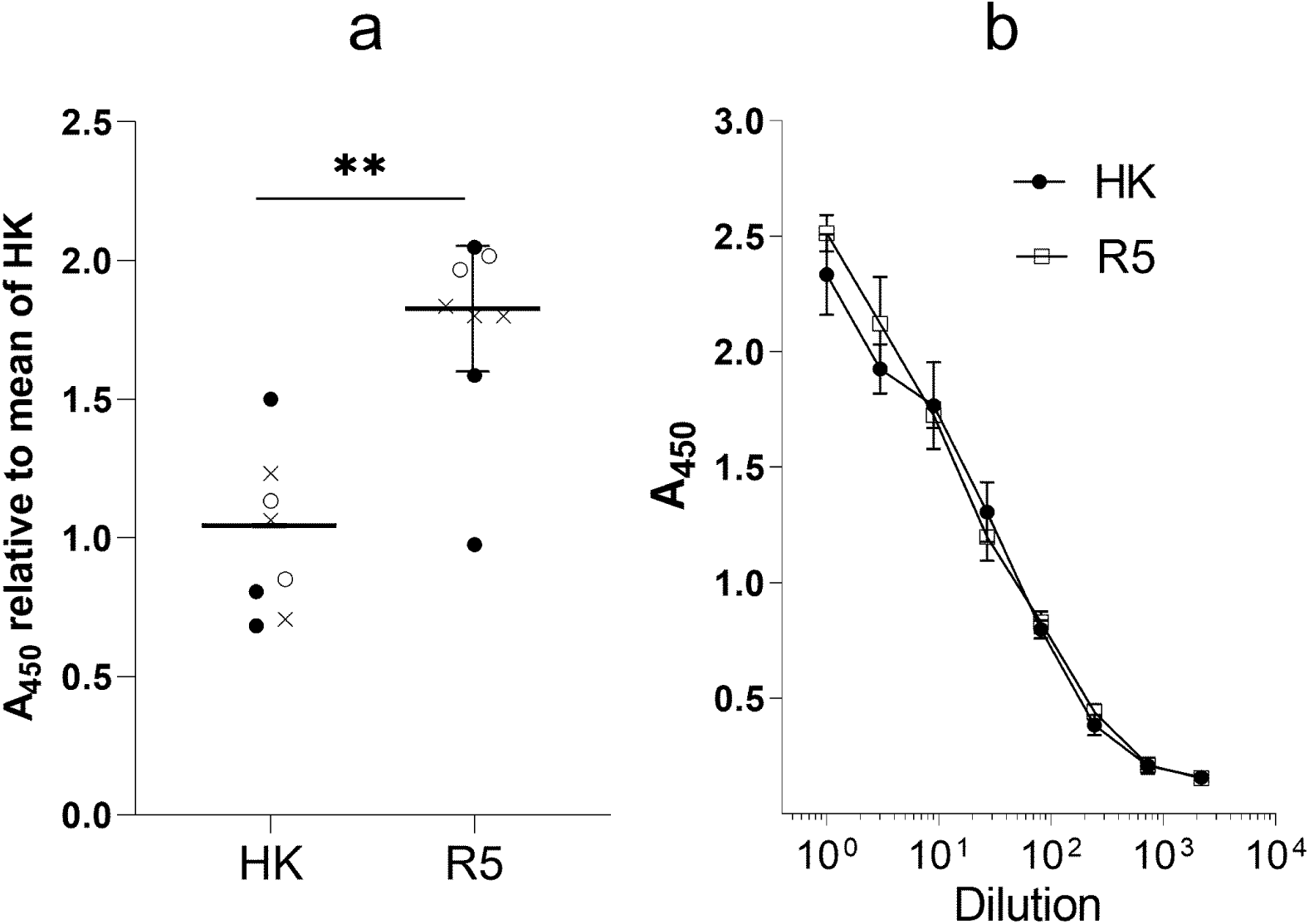
Attachment of HK and R5 to cells in HTBE cultures. (**a**) The apical sides of live HTBE cultures were washed with PBS+ to remove accumulated mucins and inoculated with 0.2 ml of DMEM-BSA containing 1.3×10^6^ FFU of HK and R5. Control cultures were inoculated with DMEM-BSA. After 1-h incubation at 4°C the cultures were washed, fixed and immune-stained using anti-HK primary antibodies and HRP-labelled secondary antibodies. The mean absorbance in the control cultures was subtracted, and the results were expressed as the relative absorbance at 450 nm (A_450_) in R5-treated and HK-treated cultures with respect to the mean absorbance in the latter. Open circles, closed circles and crosses depict individual data points from three experiments performed on different days. Mean, SD and P values were calculated using within-day averages. (**b**) Control of the concentrations of physical virus particles in suspensions of HK and R5 used for the HTBE attachment experiments. Suspensions were serially diluted in PBS and adsorbed in the wells of ELISA microplates. The wells were washed, fixed and immuno-stained as described above. Shown are the results of one experiment with 5 replicates per condition. The absorbance (A_450_) reflects non-specific binding of HK and R5 to the plastic. Overlap of the A_450_ vs dilution curves indicate that suspensions contained equal amounts of viral particles.

### 3.5. Effects of non-226/228 substitutions in the HA on virus infection in MDCK cells

In MDCK cells, R5 formed smaller plaques than did HK indicative of a less efficient multicycle replication of the former virus (Fig. 6). The effects of point substitutions in the HA on plaque size was studied separately for HK mutants and R5 mutants (Fig. 6). Although the resolving power of the assay was limited by substantial heterogeneity of the plaques formed by the same virus, we noticed reproducible effects of some of the point substitutions. Thus, HK-62, HK-81 and HK-193 formed smaller plaques in comparison with HK, whereas R5-92 and R5-144 formed smaller plaques in comparison with R5. For these five mutants, the reduced size of the plaques correlated with the elevated avidity of the virus for either 6’SLN (HK-62), MDCK cells (R5-92, R5-144), or both substrates (HK-81, HK-193) (compare Fig. 6 and Fig. 4). We previously studied dependence of replication efficiency of IAVs on their binding avidity and demonstrated that excessive avidity slowed down the release of viral progeny from infected cells and spread of the infection (Gambaryan et al., 1998b). We assume that the same mechanism explains, at least in part, the smaller plaque size of the mutants of HK and R5 containing avidity-enhancing substitutions.

**Fig. 6.**
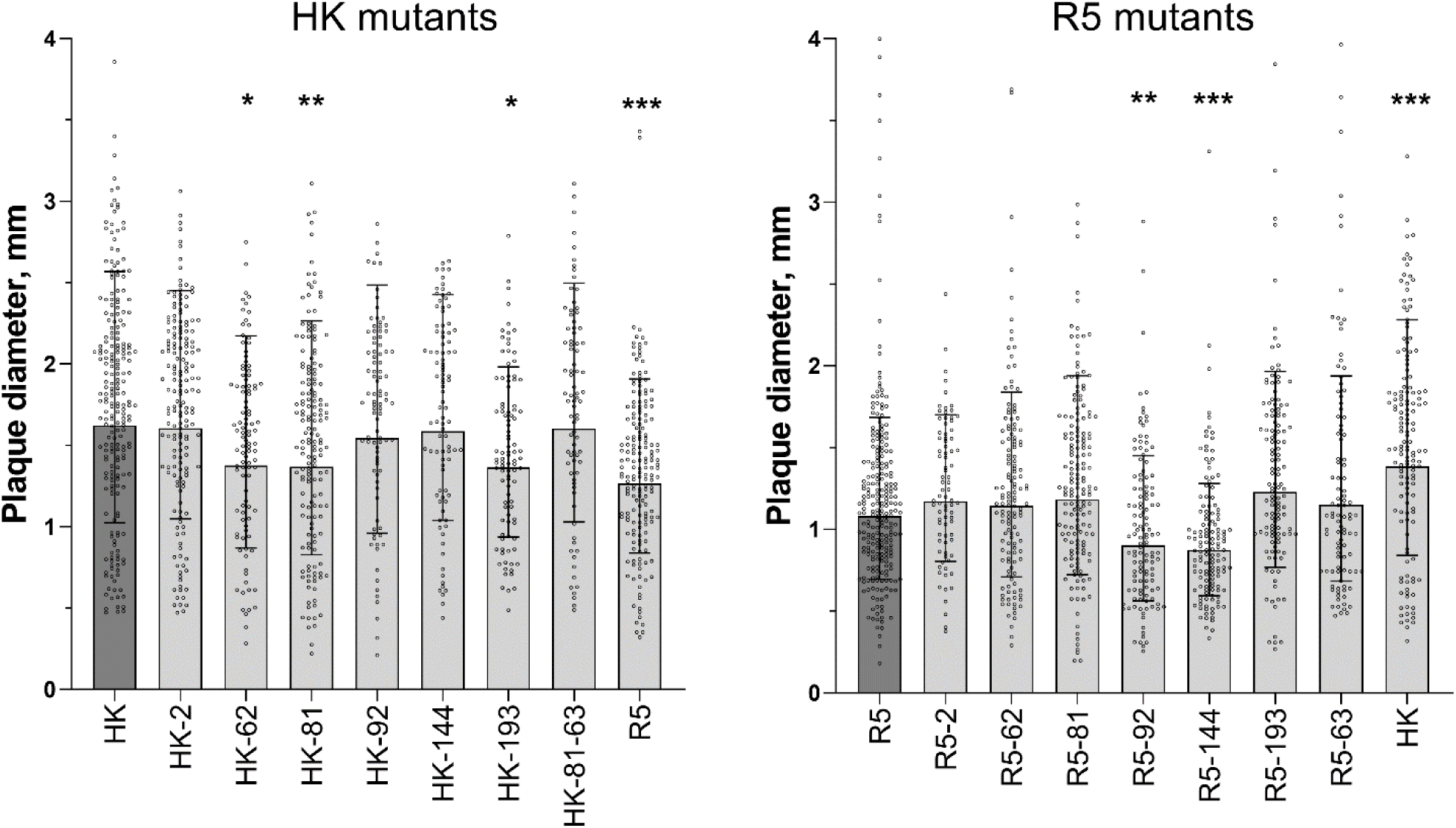
Diameter of plaques formed by viruses in MDCK cells. Cells in six-well plates were inoculated, incubated under semi-solid overlay medium for 48 h at 37°C, fixed and immunostained. Two panels represent two groups of viruses tested separately. Each panel shows diameters of individual plaques adjusted for day as described in Materials and Methods, geometric mean (bars) and geometric SDs from 1 to 4 experiments performed on different days. Asterisks depict P values for the differences between the mutants and the corresponding parental virus (HK in the left panel and R5 in the right panel).

### 3.6. Effects of non-226/228 substitutions on virus infection in HTBE cultures

Paulson and colleagues postulated that changes in receptor-binding properties of avian IAVs during their adaptation to humans may serve to increase virus binding to and infection of human airway epithelial cells and to minimize its binding to and neutralization by respiratory mucus (Baum and Paulson, 1990; Couceiro et al., 1993). To determine whether non-226/228 substitutions in the HA contributed to these effects, we inoculated differentiated HTBE cultures with either R5 or HK and determined the numbers of infected cells 8 h post-infection. To focus on the role of virus interaction with receptors on cells, the cultures were extensively washed prior to infection to remove accumulated mucins. In parallel, replicate cultures were infected in the presence of the endogenous mucins. Less cells were infected with HK than with R5 in the mucus-deprived cultures (Fig. 7a), this effect agreed with the higher binding avidity of R5 for receptor analogues and HTBE cells (Figs. 4 and 5). As expected, the presence of HTBE mucins reduced infectivity of both viruses, with HK still infecting less cells than R5. The mean percentages of cells infected in the presence of mucins with respect to the infection without mucins were 15.1% for HK and 17.2% for R5, and the difference was not statistically significant. Thus, our results suggested that substitutions separating HK from R5 reduced efficiency of entry of HK into human airway epithelial cells and that these substitutions did not make HK less sensitive than R5 to neutralization by human airway mucins.

**Fig. 7.**
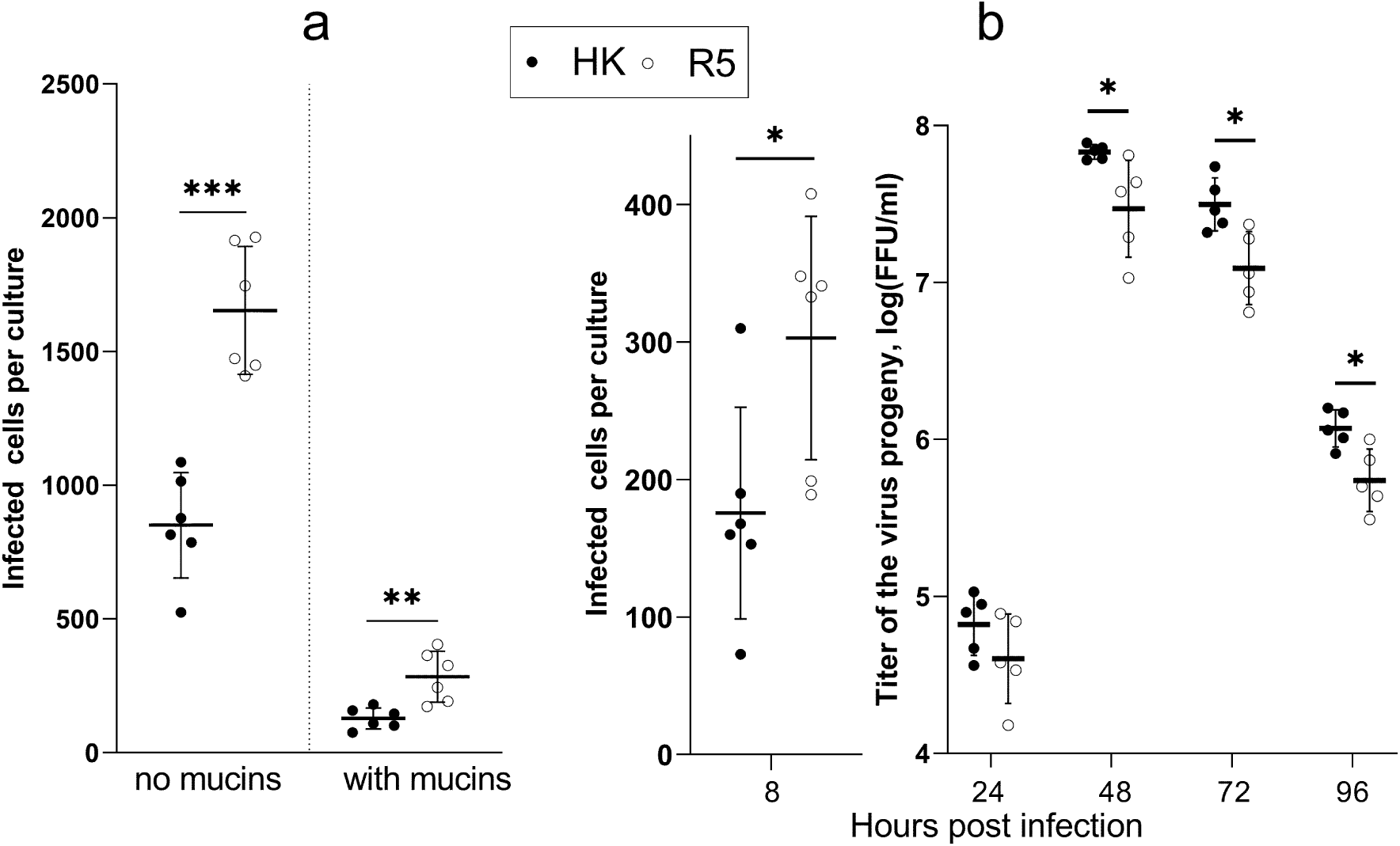
Infectivity and multicycle replication of HK and R5 in HTBE cultures. (**a**) The apical sides of HTBE cultures were washed with PBS+ to remove mucins and inoculated with 2×10^4^ FFU of HK (closed circles) and R5 (open circles) with and without addition of mucins using 6 replicate cultures per condition. The inoculum was removed after 1 h. The cultures were incubated for 7 h under ALI conditions, fixed, immuno-stained for viral NP, and numbers of infected cells were counted. (**b**) The apical sides of washed HTBE cultures were inoculated with 7×10^4^ FFU of the viruses without addition of mucins and processed as described above. Six cultures per virus were fixed 8 h post infection for immuno-staining and counting of infected cells. Viral progeny was periodically harvested by washing the apical sides of the remaining cultures, and the harvests were titrated simultaneously at the end of the experiment. P values were determined using Student’s t-test.

Reduced infectivity of HK compared to R5 in HTBE cultures was in apparent inconsistency with our previous observation of more efficient multicycle replication of HK in this cell system (Van Poucke et al., 2015). We therefore compared single- and multicycle replication of HK and R5 in the same experiment (Fig. 7b). This experiment confirmed that R5 infected more cells in the first round of infection in HTBE cultures, whereas HK produced more viral progeny after multiple infection cycles. To infer which of the substitutions separating HK from R5 contributed to more efficient multicycle replication of HK in HTBE cultures, we compared replication of HK and its single-point mutants under competitive conditions. In the first experiment (Fig. 8), we focused on substitutions at positions 81 and 193 as they showed the major effect on receptor-binding properties of HK and R5. We also studied the substitution at position 63, because it affected HA glycosylation, showed the same phenotypic effect as did substitution 81 and could have served the same function in Memphis/1968-like IAVs as did substitution 81 in Hong Kong/1968 IAVs. HTBE cultures were inoculated with 1:1 mixtures of the IAVs differing by single substitutions based on viral infectious titers in MDCK cells. The viral progeny was collected daily from the apical sides of the cultures, and compositions of the original inoculum and the harvests from day 2 and day 4 were determined by Sanger sequencing. Replication of the mixture of HK with HK-81 resulted in the enrichment of viral progeny with HK (Fig. 8a). Accordingly, replication of the mixture of HK-81-63 with its avian-type precursor HK-81 was enriched with the former virus (Fig. 8c). These results indicated that the human-type amino acid substitutions D81N, D63N and/or accompanying addition of N-glycan increased virus fitness in HTBE cultures. No significant changes in the composition of the HK mixture with the HK-193 mutant was observed after its replication (Fig. 8b) indicating that substitution at position 193 had no detectable effect on viral fitness.

**Fig.8.**
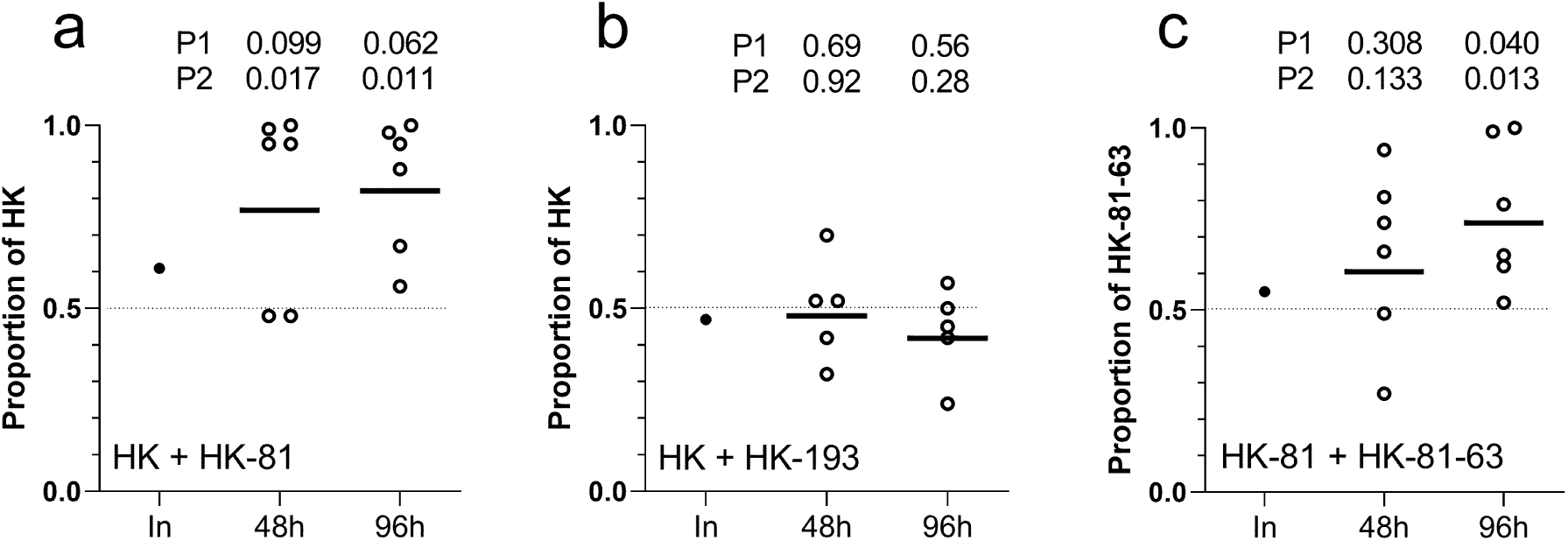
Comparison of viral fitness in HTBE cultures in competitive replication experiments. Replicate cultures were inoculated with one-to-one mixtures of HK and HK-81 (**a**), HK and HK-193 (**b**), and HK-81 and HK-81-63 (**c**) using 4 x 10^3^ FFU of each virus per culture. Proportions of viral genotypes in the original inoculum (In) and in the material harvested 48 h and 96 h post inoculation were determined by Sanger sequencing. Figures show proportion of the HA genotype of the indicated virus in the inoculum (solid circle) and in the replicate infected cultures (empty circles); solid lines depict mean values. P values refer to the differences with respect to the genotype proportion in the inoculum (P1) and with respect to the targeted 1:1 proportion in the inoculum of infectious virus particles (dotted line) (P2).

We next studied simultaneous competition of seven IAVs, HK and its six single-point HA mutants. Equivalent amounts of plaque-forming units of the viruses were mixed, and three different dilutions of this mixture were inoculated into the HTBE cultures. The first group of cultures (group L) received 5 PFU of each of the seven viruses per culture, two other groups received 20 and 320 PFU per culture (groups M and H, respectively). The viral progeny was harvested 3 days post inoculation, and the proportions of the HA gene segments with avian-type substitutions were analyzed in the harvests by next generation sequencing (Fig. 9). HA segments of all 6 mutants were present in the harvests from all cultures in the group H. By contrast, the harvests in the group M and, especially, group L were highly heterogeneous, with proportions of the mutants varying between 0.00 and 1.00. The proportions of three mutants, HK-62, HK-81, and HK-144 were significantly reduced in the harvests L as compared to their proportions in the inoculated mixture. The mutants HK-81 and HK-144 also displayed reduced frequencies in the group H. No statistically significant changes with respect to the inoculum were observed in the case of HK-92 and HK-193. In contrast with other mutants, the proportion of HK-2 was higher in the harvests than in the inoculum for all three infection doses used. These results indicated that i) HK-2 replicates more efficiently than the other mutants, ii) HK-62, HK-81, and HK-144 replicate less efficiently than the other mutants, and iii) HK-92 and HK-193 have intermediate replication efficiency.

**Fig 9.**
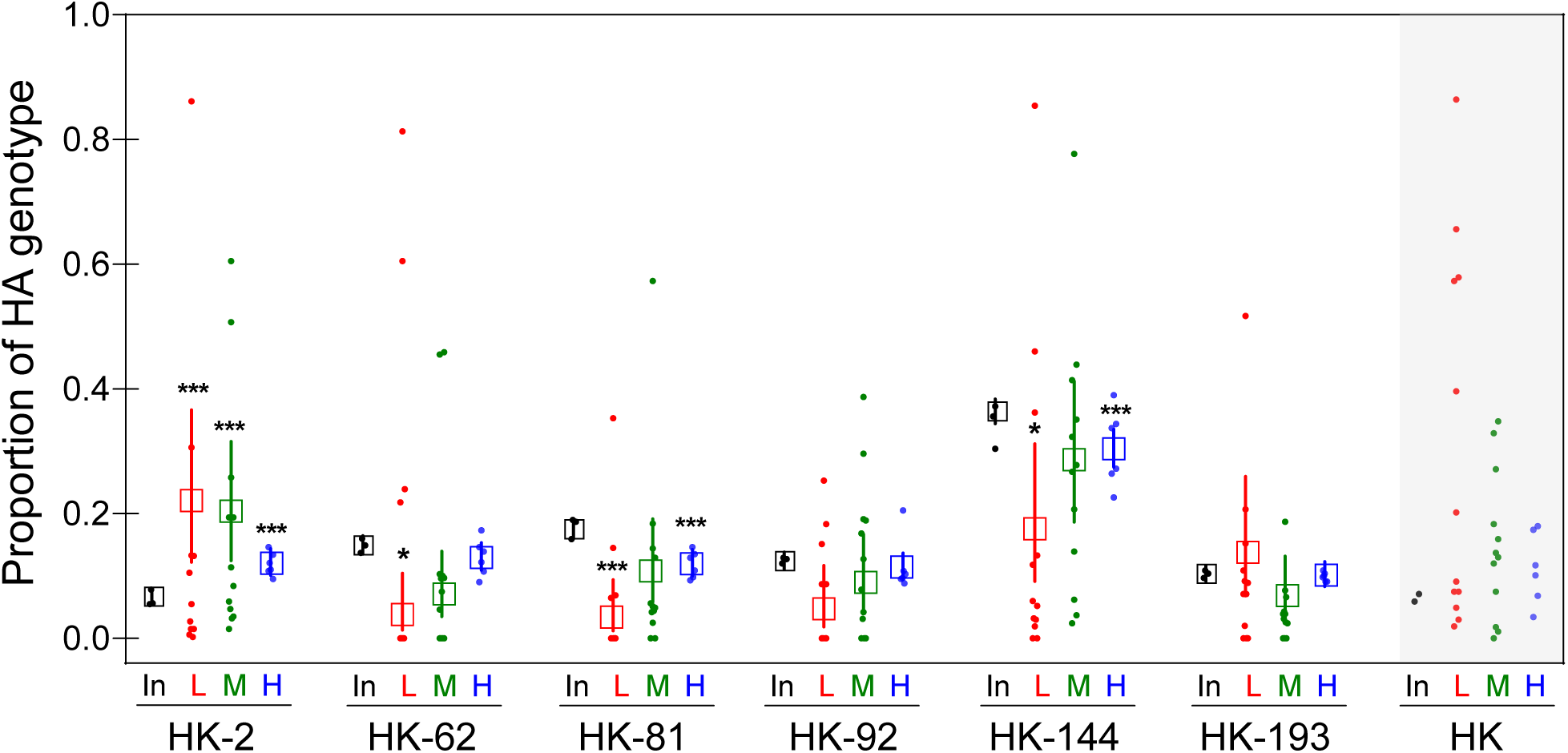
Competitive replication of HK and its 6 single-point HA mutants in HTBE cultures. HTBE cultures were inoculated with the mixtures of HK and its 6 HA mutants containing 5 PFU (L, 12 replicate cultures), 20 PFU (M, 12 replicates) and 320 PFU (H, 6 replicates) of each virus. After 1-h incubation, the inoculum was removed, the cultures were incubated under ALI conditions, and the apical material was harvested at 72 h post-inoculation. Small circles show proportions of each HA genotype determined by next generation sequencing in the inoculated mixture (In) and in each replicate harvest in the L, M and H groups. Empty squares and error bars represent the predicted mean values and confidence intervals inferred by a generalized linear model, considering the over-dispersion of the data. Asterisks depict differences between proportion of the corresponding genotype in the harvest and the inoculum. The proportions of the parent HK virus (gray background) were inferred by subtracting proportions of six HA variants from 1. These data are shown for illustration only; no hypotheses were tested.

Whereas the mutated genotypes of HK were unambiguously identified in the mixtures via unique nucleotide substitutions, the amount of the parent HK could not be directly quantified by sequencing. We therefore inferred proportions of HK in the mixtures by subtracting proportions of the mutants from the theoretical value of 1. Because of the intrinsic high errors of this approach, the data could not be analyzed statistically. One can see, however, that the proportion of HK in the harvests increased with respect to the inoculum and that the frequency in the group L was higher than in the group H. Obviously, a higher number of viral replication cycles in group L than in groups M and H was responsible for a higher enrichment of the mixture by the best-fit virus. Remarkably, this pattern of HK resembled the pattern shown by HK-2 and differed from the patterns displayed by 5 other mutants. In this view and taking into account that HK-2 mutant did not differ from HK in most phenotypic assays, we assume that HK-2 has the same fitness as does HK. Collectively, we conclude from the replication experiments in HTBE cultures that avian-type substitutions at positions 62, 63, 81 and 144 decrease in-vitro fitness of the pandemic virus, substitution at position −2 has no effect on fitness, and that effects of substitutions 92 and 193 (if any) were not statistically significant under assay conditions.

### 3.7. Airborne transmission of HK and R5 in ferrets

Transmission through the air is essential for the pandemic spread of IAVs in humans. To evaluate effects of non-226/228 substitutions in the HA of the 1968 pandemic viruses on transmissibility, we employed the ferret airborne transmission model. To avoid potential undesirable effects of the PR8-derived gene segments on viral fitness in ferrets, recombinant HK and R5 containing all 8 gene segments of A/Hong Kong/1/1968 were used in the transmission experiments. All directly inoculated ferrets shed the viruses from the nose and throat starting from day 1 after infection, the duration of shedding and peak titers did not significantly differ between HK and R5 (Fig. 10). Two of the airborne contact ferrets in the HK group became infected and shed the virus in high titers starting from the day 3 after the contact, the third contact ferret produced one virus-positive swab on the day 3. All four contact animals in the HK group seroconverted. In the R5 group, only one of four contact animals shed the virus and seroconverted. We sequenced the HA of the transmitted HK and R5 viruses present in the throat swabs of all positive indirect contact animals at 3 days post exposure and found no substitutions. Although only small numbers of animals were used in the transmission studies, a careful interpretation of these results indicated that R5 transmitted less efficiently than HK, although not significant, suggesting that substitutions other than those at the RBS were required for efficient transmission of HK virus. Although it was tempting to study effects of individual substitutions separating HK and R5 on virus transmissibility, these studies were not pursued for ethical reasons given the small differences in transmissibility of parental viruses and, hence, necessity to use large groups of animals for statistically significant detection of even smaller effects.

**Fig. 10.**
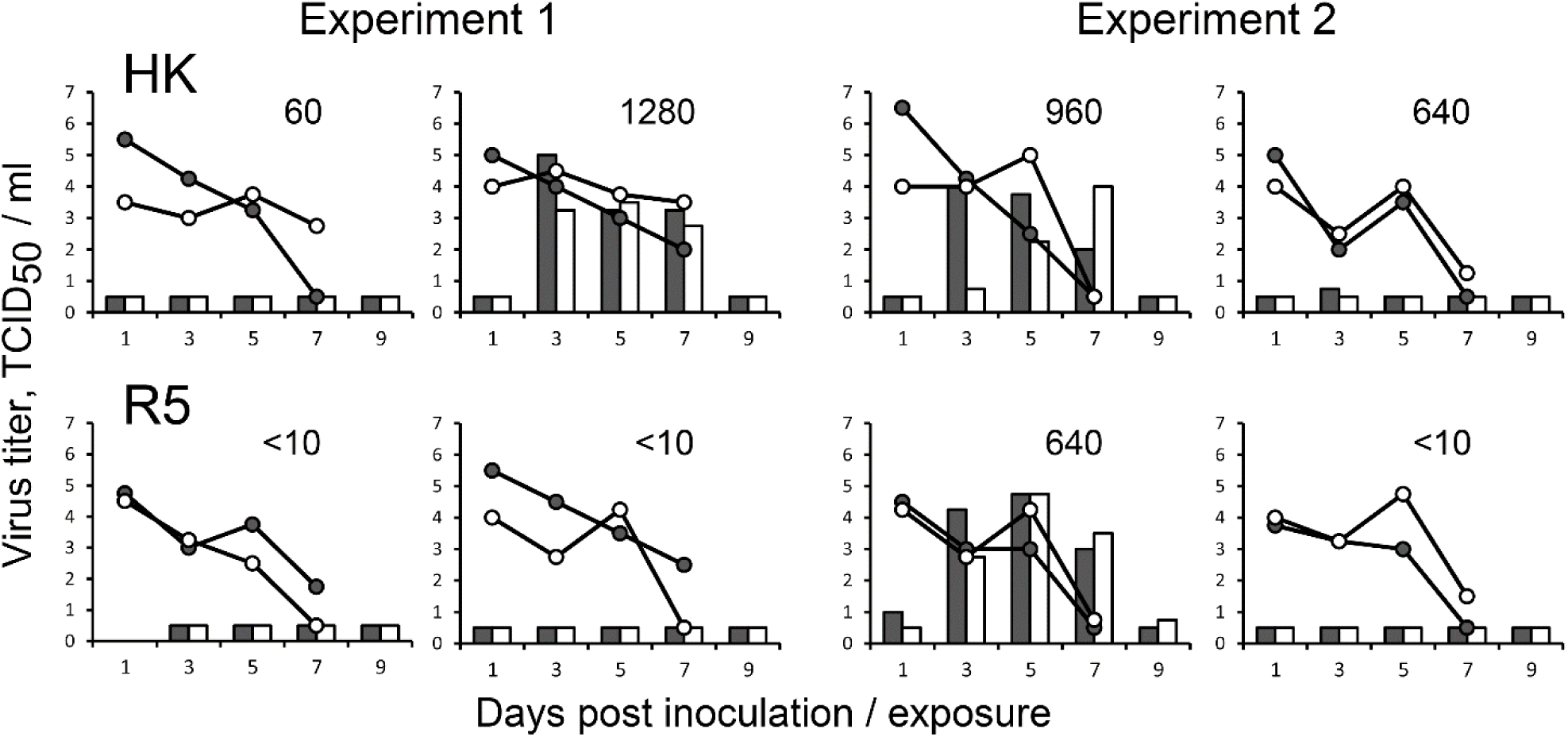
Comparison of airborne transmission of HK and R5 in ferrets. Groups of two ferrets were inoculated intranasally with 10^6^ TCID_50_ of recombinant viruses HK (top panels) and R5 (bottom panels) containing all eight gene segments of A/Hong Kong/1/1968. One naïve ferret was co-housed with each inoculated ferret in a separate transmission cage starting from one day after inoculation. Data show results of two replicate experiments performed on different days. Lines depict viral titers in nasal swabs (empty circles) and throat swabs (closed circles) collected from inoculated ferrets. White and black bars depict viral titers in nasal and throat swabs, respectively, of the indirect contact ferrets. Numbers show titers of hemagglutination inhibiting antibodies in the blood collected from the indirect contact animals, 2 weeks post exposure.

### 3.8. Analysis of H3 HA sequences in different viral host species

The H3 HA sequences available from the GISAID EpiFlu database include sequences of avian IAVs (predominantly from wild aquatic birds) and of several stable mammalian-adapted viral lineages that emerged from the avian reservoir via interspecies transmission (Fig. 11a, supplementary Fig. S5). Among them, the H3N8 equine IAVs were first recognized in 1963 (Webster et al., 1992); they continuously circulate and evolve in horses until now. The IAVs of an independent H3N8 equine lineage caused epizootic in China in 1989, circulated in horses for a few years and became extinct (Guo et al., 1995). Only one virus of this lineage, A/equine/Jilin/1/1989, was sequenced. The first of two canine H3 lineages originated from a contemporary equine H3N8 virus in Florida around 2003; the second canine lineage (H3N2) emerged from an avian precursor in Asia and was first recognized in 2006 [for reviews, see (Parrish et al., 2015; Yoon et al., 2014)]. In addition to stable mammalian-adapted lineages, the database contains a small number of IAVs isolated from mammals which cluster with avian viruses and do not form persistent mammalian lineages (supplementary Fig. S5). The human H3N2 lineage includes H3N2/1968 pandemic viruses and their permanently evolving descendants that cause seasonal influenza epidemics. Finally, multiple lineages of H3N2 swine IAVs all originated from independent human-to-swine transmissions of seasonal IAVs followed by evolution in pigs (Anderson et al., 2020).

**Fig.11.**
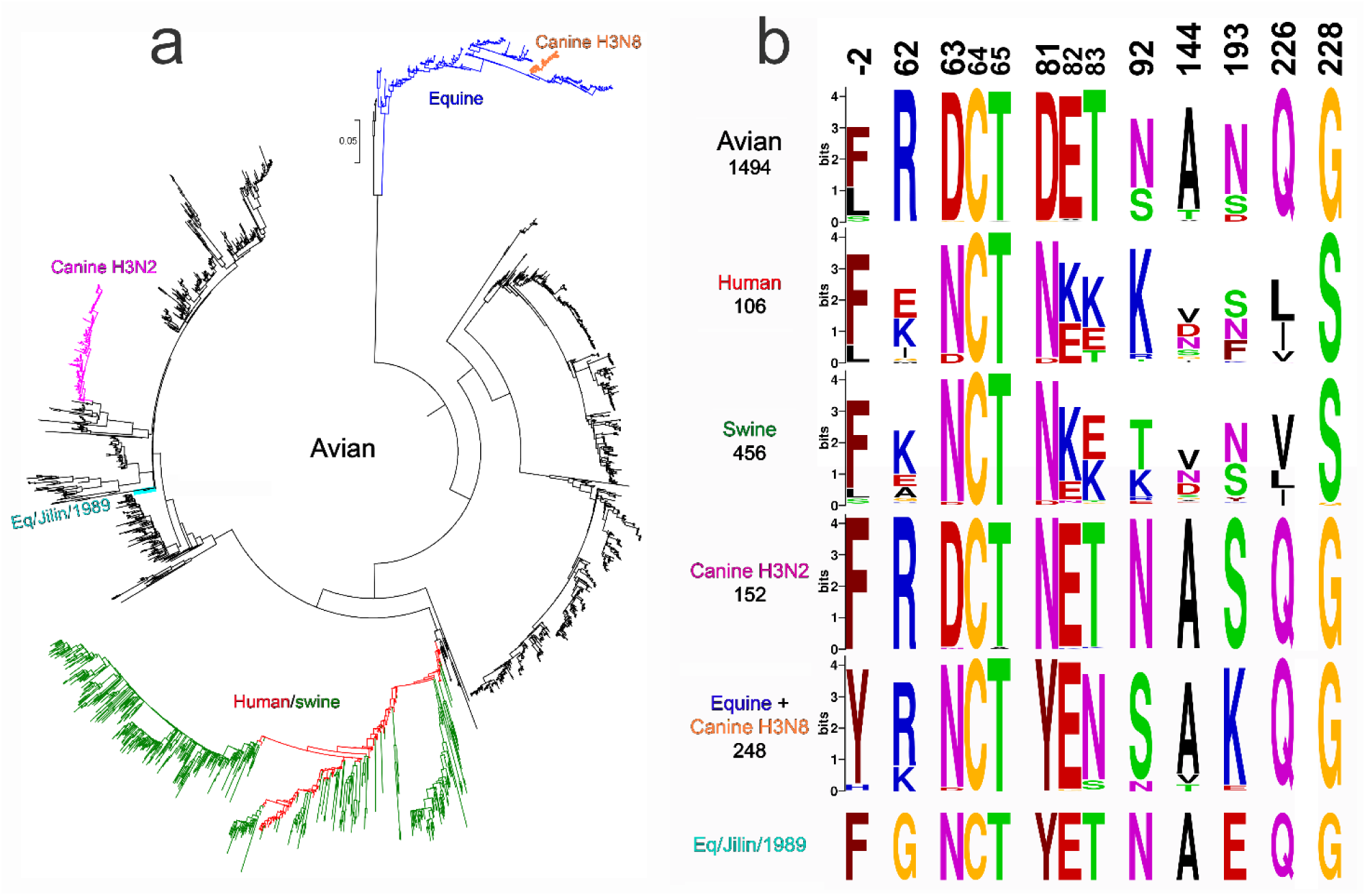
Host-specific lineages of IAVs with H3 HA and variation of amino acids at selected HA positions. (**a**) Phylogenetic tree for the H3 HA nucleotide sequences of representative sequences of human and swine viruses and all unique sequences of other mammalian and avian viruses available from GISAID EpiFlu database. The numbers of analysed sequences are shown in panel b below the lineage name. Supplementary figure S5 shows the same tree with strain names, accession numbers and amino acids at 9 HA positions under study. (**b)** Protein logos for indicated HA positions of the viral lineages shown in panel a. The overall height of each stack of letters depicts sequence conservation measured in bits. The height of each letter is proportional to the frequency of the corresponding amino acid in the alignment, the letters are ordered from most to least frequent. Only one sequence was available for the Eq/Jilin/1989 lineage.

We analysed prevalence of specific amino acids at nine HA positions in question in avian IAVs and IAVs of different mammalian lineages (Fig. 11b, supplementary Tab. S1, supplementary Figs. S5 and S6). We also estimated selective pressures on these positions in different hosts using codon-based likelihood methods FEL, Contrast-FEL and MEME, which compare rates of synonymous and non-synonymous substitutions (Kosakovsky Pond and Frost, 2005; Kosakovsky Pond et al., 2021; Murrell et al., 2012), and DEPS/FADE to identify directional evolution along human internal branches (Kosakovsky Pond et al., 2008) (Tab.1). Finally, we compared HA sequences of the earliest equine and canine isolates with sequences of their closest avian counterparts to identify potential amino acid substitutions at 9 HA positions and their correlation with substitutions in the H3N2/1968 pandemic viruses (Tab. 2, supplementary Fig. S5). Human-origin swine viruses were not included in the latter two analyses as we were interested in the avian-to-mammalian shifts. The results of these studies are summarized below.

**Table 1.** Selection pressure analysis of 9 sites of the HA of avian, mammalian and human IAVs ^a^.

| Site | Substitutions (Syn : Non-Syn) <sup>b</sup> |  |  | Pervasive selection<br>dN/dS, FEL significance for dN≠dS <sup>c</sup> |  |  | Episodic positive selection number<br>of branches, MEME significance <sup>d</sup> |  |  | Directional selection<br>(target in H, empirical<br>Bayes factor) <sup>e</sup> | Comparative selection <sup>f</sup> |
| --- | --- | --- | --- | --- | --- | --- | --- | --- | --- | --- | --- |
|  | A | M | H | A | M | H | A | M | H |  |  |
| -2 | 13:14 | 0:1 | 2:5 | 0.491 * | 0.354 | 1.89 | 0 | 0 | 0 |  |  |
| 62 | 18:1 | 1:2 | 2:5 | 0.0237 *** | 0.548 | 1.57 | 0 | 0 | 0 |  | H>A ***, overall *** |
| 63 | 4:5 | 1:2 | 1:2 | 0.145 *** | 0.767 | 0.782 | 0 | 0 | 0 | N (37) |  |
| 81 | 3:7 | 0:0 | 0:1 | 0.378 * | 0 | 0.675 | 5 | 0 | 0 |  | overall * |
| 92 | 10:16 | 1:2 | 0:2 | 0.504 * | 0.616 | 0.734 | 0 | 0 | 0 | T (41) |  |
| 144 | 13:24 | 0:3 | 1:8 | 0.807 | 1.05 | 3.04 * | 0 | 0 | 0 * |  | H>A * |
| 193 | 16:42 | 0:3 | 0:7 | 0.679 | 0.464 | 1.01 | 0 | 3 | 0 |  |  |
| 226 | 14:0 | 0:0 | 3:15 | 0 *** | 0 * | 6.98 *** | 0 | 0 | 7 *** |  | H>A ***, H>M ***, overall *** |
| 228 | 17:0 | 1:1 | 2:0 | 0 *** | 0.23 | 0 ** | 0 | 0 | 0 |  |  |

<sup>a</sup> Groups A, M and H contained, respectively, 1492 sequences of avian IAVs, 406 sequences of equine, canine, feline and seal IAVs and 803 sequences of human IAVs isolated in the years from 1968 to the end of 1999 (set H). We restricted analyses only to internal branches (see Materials and Methods). Asterisks depict significance levels for the tests as follows: \*, $P < 0.05$ ; \*\*, $p < 0.01$ ; \*\*\*, $p < 0.001$ .
<sup>b</sup> The number of synonymous and non-synonymous substitutions inferred to have occurred on internal branches in the corresponding group set with the SLAC method.
<sup>c</sup> Site-level dN/dS estimate along internal branches in the corresponding group inferred using the FEL method. Asterisks show significance levels for the test that $dN \neq dS$ . When significant result is obtained, negative selection is inferred if $dN/dS < 1$ , otherwise diversifying positive selection is inferred.
<sup>d</sup> Site-level characterization of episodic positive selection in the corresponding group inferred using the MEME method. Asterisks show significance levels for the test that $dN > dS$ for some fraction of branches. The number of individual branches where episodic selection may have acted is indicated as well (empirical Bayes factor $\geq 100$ )
<sup>e</sup> Site-level characterization of directional selection using the DEPS/FADE method. The residue that is the putative target of directional selection is indicated, with empirical Bayes factor supporting directional selection shown.
<sup>f</sup> Site-level characterization of differences in selective pressures between branch groups using the Contrast-FEL method. Asterisks show significance levels for the test that dN/dS differ between pairs of branch groups, or among all branch groups (overall). Notation like H>A means that dN/dS for “human” branches is greater than dN/dS for “avian” branches. Only significant results are listed.

**Table 2.** Alteration of amino acids at 9 positions of the H3 HA during host shifts ^a^.

| Host-specific virus lineage<br>and the earliest isolate | Position |  |  |  |  |  |  |  |  |
| --- | --- | --- | --- | --- | --- | --- | --- | --- | --- |
|  | -2 | 62 | 63 | 81 | 92 | 144 | 193 | 226 | 228 |
| **Human/swine**<br>Hong Kong/1/1968<br>Memphis/1/1968 | F→L | R→I | D→N* | D→N* | N→K | A→G | N→S | Q→L | G→S |
| **Equine**<br>Equine/Miami/1/1963 (H3N8) | F/L/Y→H |  | D→N* | D→Y |  |  | N→K |  |  |
| **Canine H3N2**<br>Canine/Guangdong/1/2006 |  |  |  | D→N* |  |  |  |  |  |
| **Eq/Jilin/1989**<br>Equine/Jilin/1/1989 (H3N8) |  | R→G | D→N* | D→Y |  |  | N/D→E |  |  |
| **Sporadic isolates of avian-like IAVs from mammals <sup>b</sup>** |  |  |  |  | N→S |  | N→S |  |  |

<sup>a</sup> Table shows substitutions at the indicated positions that separate the earliest isolates of stable mammalian lineages from the phylogenetically closest avian viruses. Empty cells indicate a lack of changes. Asterisk next to N indicates that substitution generates glycosylation site. The analysis was performed using 2489 HA sequences depicted in the Fig.11 and in supplementary Fig. S5.
<sup>b</sup> The HAs of 10 swine, canine and seal IAVs which clustered with HAs of avian viruses but did not form stable lineages. Substitutions were observed in two viruses, A/harbour seal/New Hampshire/179629/2011 (H3N8) (N92S) and A/seal/Massachusetts/3911/1992 (H3N3) (N193S)

#### Position −2

FEL predicts that this position is under pervasive negative selection (dN/dS = 0.49) in avian IAVs; the site exhibits a large number of substitutions (both synonymous and non-synonymous) on avian branches (see supplementary Fig S6). No significant selection effects were detected by other likelihood methods (Tab. 1). Amino acids F and L are typically present at this position in the HAs of avian, human and swine IAVs (Fig. 11b, supplementary Figs. S5 and S6). The earliest isolates of the A/equine/Miami/1963-like (H3N8) IAVs had H in position −2, whereas closest avian IAVs carry F, L and Y (Tab. 2, supplementary Figs. S5 and S6). However, given a significant divergence of this equine lineage from other IAVs, the identities of amino acids at HA position −2 of an avian precursor and first equine-adapted variants remain unclear. No changes at this position occurred during emergence of other mammalian lineages. Collectively, these analyses provided no indications of the association of the substitution F(−2)L with viral host range and interspecies transmission.

#### Position 62

The codon is negatively selected in birds overall (FEL) with 99.3% of analysed avian HAs containing R_62_ (supplementary Tab. S1). Contrast-FEL detects a higher dN/dS in human IAVs compared to avian IAVs; the point estimate of dN/dS in humans is 1.57, but this is not significantly different from 1. This finding is consistent with the location of the amino acid in the antibody-binding site E of the H3 HA (Wiley and Skehel, 1987) and its evolution under immune selection pressure in humans. In fact, all positions analysed here, with the exclusion of position −2, are also located in the antibody-binding sites A (residue 144), B (193), D (226, 228) and E (62,63,81,92). Avian-type R_62_ is conserved among canine and most equine IAVs, with conservative substitution R to K in equine viruses isolated after 2008 (Fig. 11b, supplementary Figs. S5 and S6). The H3N2/1968 pandemic viruses acquired non-conservative substitution R62I. Another independent host switch event, transmission of an avian IAV to horses in Asia in 1989, was also accompanied by non-conservative substitution R62G (Tab. 2). Two independent non-conservative substitutions of conserved avian-type residue R_62_ suggest potential adaptive role of these substitutions during avian-to-mammalian transmission.

#### Positions 63 and 81

Avian HAs contained D_63_ and D_81_ in 98.3% and 98.6% of analysed sequences, respectively (Fig. 11b, supplementary Tab. S1). Both codons are negatively selected in birds overall (FEL). The substitution from D to N at either position 63 or position 81 of two co-circulating pandemic virus lineages generated glycosylation sites. Remarkably, the substitution D63N and acquisition of glycosylation site accompanied emergence of both equine IAV lineages, whereas substitution D81N with new glycosylation site occurred during transmission of an avian H3N2 virus to dogs (Tab. 1). The glycosylation sites became fixed in H3N8 equine and H3N2 canine lineages, moreover, the equine-origin H3N8 canine and human-origin H3N2 swine IAVs inherited and preserved the glycosylation sites of their mammalian precursors. As a result, all known mammalian IAVs with H3 HA differ from H3 avian IAVs by the presence of N-glycan at either position 63 or position 81 (Fig. 11b, supplementary Figs. S2, S5 and S6). Observed parallel evolution of amino acids at positions 63 and 81 during avian-to-mammalian adaptation represents a strong indication of their adaptive role in H3N2/1968 pandemic IAVs. There is evidence of directional evolution towards N on the human branches at site 63. Furthermore, substitution D81Y occurred in both H3N8 equine lineages. Thus, alteration of the properties of amino acid in position 81 may play an adaptive role in interspecies transmission irrespectively from and/or in addition to the effect of the substitution on HA glycosylation.

#### Position 92

Avian IAVs contain either N_92_ or S_92_. The codon is negatively selected in birds overall (FEL, dN/dS = 0.504) (Tab. 1). Whereas none of the avian HA sequences contained K_92_, non-conservative substitution N92K altering change of the amino acid side chain accompanied emergence of the pandemic IAVs. Of note, this substitution is located in the close proximity of another charged human-type substitution R62I (see Fig. 1b) and could compensate for the effect of the latter substitution on the net surface charge of the protein in this area. No substitutions in position 92 occurred during emergence of other stable mammalian lineages, however, conservative substitution N92S was present in H3N8 IAVs that caused epizootic with cases of fatal pneumonia in New England harbour seals in 2011 (Anthony et al., 2012) (Tab.2, supplementary Fig. S5). Along the human branches, there is evidence of directional selection towards T, with two clades showing N→T substitutions which were then maintained.

#### Position 144

In accord with location of the amino acid in the antigenic site, codon 144 is under positive selection in humans (FEL dN/dS = 3.04, and MEME). dN/dS on human branches is significantly higher than on the avian branches (Contrast-FEL). Avian IAVs typically contain A, I, or V but never G. By contrast, the H3N2/1968 pandemic IAVs carried a non-conservative substitution A144G, which could affect the structure of polypeptide loop 140-145 located in the vicinity of the RBS. These notions suggest potential functional significance of this substitution for the avian-to-human adaptation. There is no evidence of parallel evolution at this site in other instances of avian-to-mammalian transmissions (Tab. 2).

#### Position 193

This amino acid is located at the upper rim of the RBS. The codon is predicted to be under negative selective pressure in horses and dogs, but not in other species (Tab. 1). Avian HAs contain N and, less frequently, S or D. Unique substitutions to K_193_ and E_193_ occurred during independent transmissions of avian precursors to horses (A/equine/Miami/1963-like lineage and A/equine/Jilin/1/1989-like lineage); conservative substitution N-to-S was found in H3N3 avian-like IAVs isolated from seals in 1992 (Tab. 2, supplementary Fig. S5). These findings suggest a potential functional role of the substitution in position 193 during interspecies transmission.

Amino acids Q_226_ and G_228_ are critical for the avian HA binding to avian-type receptor motif Neu5Acα2-3Gal (Gamblin et al., 2020; Matrosovich et al., 2006b; Shi et al., 2014). Both codons are under purifying selection in birds, horses and dogs, which share binding preference for Neu5Acα2-3Gal-terminated receptors. By contrast, the codon 226 is under pervasive and episodic positive selection in humans (FEL dN/dS = 6.98, MEME), and significantly higher dN/dS in humans compared to both avian and mammalian lineages.

## 4. Discussion

Unavailability of immediate animal precursors of pandemic IAVs hampers understanding of genetic and phenotypic changes in the HA that were essential for the viral animal-to-human adaptation and pandemic spread. To mitigate this problem, we generated and characterized the recombinant IAV R7 containing the HA of the hypothetical most recent common ancestor of H3 avian and H3N2/1968 pandemic IAVs. R7 displayed receptor-binding profile typical for duck viruses and differed in this respect from IAVs perpetuated by gulls, shorebirds and gallinaceous land-based poultry. The HA of R7 showed high conformational stability and low pH optimum of fusion compatible with the aquatic bird origin of the H3N2/1968 HA (Baumann et al., 2016; Scholtissek, 1985). These properties of R7 agreed with the hypothesis that the H3N2/1968 pandemic IAVs originated from a duck virus (Bean et al., 1992; Kida et al., 1987). IAVs with the HA sequences highly similar to R7 were isolated from both wild migratory ducks captured on a Pacific flyway in Japan and domestic ducks in Southern China (Kida et al., 1987; Yasuda et al., 1991) (supplementary Fig. S1b). It seems likely that the precursor wild duck virus was transmitted to humans via domestic ducks either with or without additional intermediate host species.

The HAs of all four characterized pandemic IAVs (H1N1/1918, H2N2/1957, H3N2/1968 and H1N1/2009) were relatively stable, whereas swine IAVs, highly pathogenic H5 and H7 IAVs from gallinaceous poultry and some IAVs of aquatic birds display low conformational stability (Baumann et al., 2016; Galloway et al., 2013; Russell et al., 2018). These observations suggested that adaptation of animal IAVs to humans may require stabilizing substitutions in the HA (Russell et al., 2018; Russier et al., 2016), however, it remained unclear whether this mechanism contributed to the emergence and initial pandemic spread of any known pandemic virus. We found that the HA of HK was in fact slightly less stable than the precursor HA of R7 with a pH_50_ of conformational transition of 5.4 and 5.25, respectively (Fig. 2). Reduced HA stability of HK was primarily associated with substitutions at positions 226 and 228; substitutions at other HA positions had lower if any effects. Thus, the duck precursor of the H3N2/1968 IAVs had a sufficiently stable HA and was able to adapt to humans without elevation of its conformational stability.

Analysis of the receptor-binding specificity of HK and its HA variants (Fig. 3b) confirmed the concept that preferential binding of the H3N2/1968 viruses to Neu5Acα2-6Gal-terminated receptors is primarily determined by substitutions Q226L and G228S (Connor et al., 1994; Matrosovich et al., 2000). The combination of other human-type substitutions in the HA decreased binding of HK to both Neu5Acα2-6Gal- and Neu5Acα2-3Gal-terminated receptor analogues, reduced its attachment to apical surfaces of HTBE cultures and lowered infectivity for HTBE cells without affecting efficiency of virus neutralization by human airway mucus (Figs. 4,5 and 7). These results indicated that non-226/228 substitutions lowered the avidity of HA binding to receptors on human target cells. Although HK infected less cells than R5 during initial inoculation into the HTBE cultures, HK produced more infectious virus particles after multicycle replication (Fig. 7), suggesting that it outperforms R5 during post-entry replication stage(s). The reduced binding avidity can increase fitness of HK by facilitating release of viral progeny from cells and preventing its receptor-mediated self-aggregation (Gambaryan et al., 1998b; Kaverin et al., 2000). In addition, some of the non-226/228 substitutions could, in principle, promote replication of HK relative to R5 by avidity-independent mechanisms, such as facilitation of synthesis and intracellular processing of the HA protein or assembly of virus particles. Further studies are needed to clarify potential roles of these mechanisms.

Our observation of reduced HA avidity of H3N2/1968 IAVs is in line with the observed lower avidity of the HA of swine-origin H1N1/2009 pandemic IAV as compared to its closest available swine counterparts (de Vries et al., 2011; Xu et al., 2012). The NA catalytic activity of H1N1/2009 was also lower than that of swine IAV NAs (Xu et al., 2012), in agreement with the concept that a functional balance between HA and NA is essential for efficient replication and transmission of IAVs (de Vries et al., 2020; Wagner et al., 2002). The H3N2/1968 pandemic virus was a reassortant containing the H3 HA of an avian parent and the N2 NA of a human parent. This NA differed from typical avian N2 NAs by substrate specificity (Baum and Paulson, 1991; Kobasa et al., 1999) and by substitutions in the second sialic acid binding site that reduced catalytic activity (Du et al., 2019; Uhlendorff et al., 2009). We speculate that the reduction of binding avidity of the avian-origin HA of the H3N2/1968 IAVs could have simplified its functional match with the human-origin NA.

Reduced avidity of the HK HA was primarily associated with the avian-to-human substitutions R62I, N193S and either D81N or D63N (Fig. 4). The substitution R62I is located relatively far from the receptor binding site and decreases the local and the net positive charges of the HA. The negative effect of this substitution on binding avidity could be partially associated with the reduction of electrostatic attraction of IAV particles to negatively charged soluble sialoglycans and cell membranes (Gambaryan et al., 1998b; Hensley et al., 2009). Amino acid 193 is located at the upper rim of the receptor-binding site. Charged substitutions at this position, such as N/S → K/D, were shown to affect receptor-binding properties of avian and equine viruses with different HA subtypes (Gambaryan et al., 2018; Gambaryan et al., 2012; Matrosovich et al., 2000; Medeiros et al., 2004; Peng et al., 2018). In the crystal structures of the avian H5 HA and canine H3 HA complexed with avian-type receptor glycans 6-Su-3’SLN and 6-Su-SLe^x^, the side chain of K_193_ interacts with the sulfogroup attached to GlcNAc-3 (Collins et al., 2014; Xiong et al., 2013). In the H3N2/1968 HA complexes with human-type receptor analogues LSTc and 6SLN-LN, the side chain of S_193_ contacts the Gal-4 residue of the glycan (Eisen et al., 1997; Wu and Wilson, 2020). These observations suggest that substitution N193S affects binding avidity of HK by altering HA interactions with sub-terminal saccharide residues of both avian-type and human-type receptor glycans. The N-glycans on the HA globular head typically decrease binding avidity with the effect being dependent on glycan structure and location with respect to the receptor-binding site [for review, see (Matrosovich et al., 2006b)]. Substitutions D81N and D63N are located in the same area of the HA and result in addition of structurally similar complex type N-linked glycans containing up to 4 antennae (An et al., 2015). We assume that bulky NG_63_ and NG_81_ either reach the lower rim of the RBS and directly interfere with HA-receptor interactions or have some yet undefined allosteric negative effect on binding.

The essential role of HA substitutions Q226L and G228S in the emergence of H2N2/1957 and H3N2/1968 pandemic IAVs is well established. To test whether other substitutions separating the HA of HK from its avian precursor were at all required for the avian-to-human adaptation, we compared transmission of HK and R5 via airborne droplets in ferrets, the currently preferred animal model for prediction of IAV replication and transmissibility in humans (Belser et al., 2018). R5 transmitted less efficiently than HK (Fig. 10) supporting the concept that some of the non-226/228 substitutions in the HA contributed to the human adaptation and pandemic spread of H3N2/1968 IAVs (Van Poucke et al., 2015). Unfortunately, the low statistical power of the current ferret transmission model with small group sizes did not allow us to study effects of individual substitutions on virus fitness and transmissibility. Additional experiments are needed to address this question, for example, analyses of virus replication and transmission in ferrets assisted by deep mutational scanning of the positions of interest (Soh et al., 2019). Alternatively, improved methods to assess virus transmissibility need to be developed.

To rank the substitutions in the order of their potential importance for the avian-to-human adaptation of the H3N2/1968 HA we took into account various adaptation-related characteristics, such as significant effect of the substitution on the HA phenotype, its location in the functional region of the HA, and dissimilar patterns of evolution of corresponding positions in birds and mammals (Tab. 3).

**Table 3.** Characteristics of avian-to-human amino acid substitutions in pandemic H3N2/1968 viruses.

|  | **F(-2)L** | **R62I** | **D63N** | **D81N** | **N92K** | **A144G** | **N193S** | **226+228** |
| --- | --- | --- | --- | --- | --- | --- | --- | --- |
| **Phenotypical effects <sup>a</sup>** |  |  |  |  |  |  |  |  |
| Preference for Neu5Ac2-6Gal-terminated receptors |  |  |  |  |  |  |  | + |
| Binding avidity |  | + | + | + | + | + | + | + |
| pH of conformational transition |  |  |  |  |  |  |  | + |
| Fusion activity (polykarion/NH<sub>4</sub>Cl) | + |  | + | + |  |  |  | + |
| Replication in MDCK |  | + |  | + | + | + | + | + <sup>b</sup> |
| Replication in HTBE |  | + | + | + |  | + |  | + <sup>b</sup> |
| **Structural features** |  |  |  |  |  |  |  |  |
| Location in the functional region of the HA | + |  |  |  |  | + | + | + |
| Alteration of charge/non-conservative substitution |  | + | + | + | + | + |  | + |
| Addition of N-glycan |  |  | + | + |  |  |  |  |
| **Variation of the codon in H3 HAs** |  |  |  |  |  |  |  |  |
| Purifying selection in avian IAVs | + | + | + | + | + |  |  | + |
| Conserved amino acid in avian IAVs |  | + | + | + |  |  |  | + |
| Human-type amino acid is not found in avian IAVs |  | + | + | + | + | + |  | + |
| Parallel evolution in mammalian IAVs |  | + | + | + |  |  | + | + <sup>b</sup> |
| **Total <sup>c</sup>** | **3** | **8** | **9** | **10** | **5** | **6** | **4** | **12** |

<sup>a</sup> Plus indicates that point substitution in corresponding position affects properties of either HK, R5 or both viruses in one or more phenotypical assays used.
<sup>b</sup> These characteristics were described in the literature (Bateman et al., 2008; Connor et al., 1994; Matrosovich et al., 2007)
<sup>c</sup> Total number of plusses in the column. This number reflects probability of the adaptive role of the substitution during the avian-to-human transmission of the HA.

The substitutions Q226L/G228S displayed the maximal total score in such combined analysis, supporting the validity of this approach. Among the non-226/228 substitutions, D63N and D81N showed the highest score. Moreover, these HA positions were characterized by a remarkable parallel evolution during interspecies transmission events (Fig. 11, Tab. 2, supplementary Figs. S5,S6). In the context of HK HA, either substitution reduced the binding avidity and slightly elevated the pH optimum of viral fusion within endosomes (Fig. 4, supplementary Fig. S3). However, it seems unlikely that these effects alone could explain addition of a novel N-glycan in all independent cases of avian H3 HA adaptation to such distinctive hosts as humans, dogs and horses. It is also unlikely that NG_63_/NG_81_ served to mask HA antigenic epitopes, given a lack of herd immunity in mammals during emergence and initial epidemic spread of a novel IAV. The HA of the HK-like strains A/X31 and A/Aichi/2/1968 containing NG_81_ was often used as a model in the general research on the role of N-glycans in protein folding, quality control and intracellular transport (Daniels et al., 2003; Gallagher et al., 1992; Hebert et al., 1997). These studies showed that folding, formation of disulfide bonds and quality control of the nascent HA chain in the ER is largely regulated by concerted interactions of N-glycans attached in critical HA regions with lectin chaperones calnexin and calreticulin. NG_81_ was found to engage calreticulin, and point mutants lacking NG_81_ displayed delay in folding due to a less efficient formation of the critical intrachain disulfide bond C64-C76 located in vicinity of this glycan (Daniels et al., 2003; Hebert et al., 1997). We therefore hypothesize that the addition of either NG_81_ or the structurally equivalent NG_63_ increases fitness of avian-origin HA in mammals by ensuring its interactions with mammalian chaperones. Of note, two other pandemic viruses, H1N1 from 1918 and 2009, contained NG_94_ in the same area of the HA. Although fitness-enhancing mechanisms of substitutions D63N/D81N remain to be fully characterized, we conclude that these substitutions represent a previously unrecognized important marker of avian-to-mammalian adaptation and pandemic potential of IAVs.

The relative importance of the other substitutions for the adaptation is less clear. R62I shows a high score (Tab. 3) and represents particular interest because it reduces HA avidity, increases viral replicative fitness in MDCK cells and HTBE cultures and because codon 62 displays distinctive evolution in avian and mammalian IAVs. Substitutions N193S and A144G are located in the functionally important region on the opposite rims of the RBS, show phenotypes in receptor-binding assays and could serve to fine-tune HA interactions with receptors in humans. The substitution F(−2)L in the signal peptide was neutral in most phenotypic and genotypic analyses performed, however, this substitution showed weak effect on viral membrane fusion activity, thus deserving attention in the future studies.

The research on mammalian adaptation of avian IAVs was strongly stimulated and advanced by two independent reports on HA substitutions that allowed airborne transmission of avian H5N1 IAVs in ferrets (Herfst et al., 2012; Imai et al., 2012). In each study, two substitutions in the RBS changed HA binding preference from Neu5Acα2-3Gal motif to Neu5Acα2-6Gal motif, one substitution removed N-glycan from the tip of the HA thus increasing binding avidity and one substitution increased HA conformational stability. Remarkably, apart from the alteration of the Neu5Ac-Gal linkage specificity, other ferret-adaptation changes in the H5N1 HA (alterations of HA avidity, stability and N-glycosylation) are discordant with the changes that accompanied emergence of the pandemic H3N2/1968. This discrepancy can be explained, at least in part, by the differences between properties of poultry-adapted H5N1 IAVs and duck-origin precursor of H3N2/1968 and between factors required for airborne transmission in ferrets and pandemic spread in humans. In any case, our results highlight the importance of the studies on previous pandemic IAVs for the influenza risk assessment and preparedness.

## Supplementary materials

**Supplementary Method.** Construction of a GAMLSS model to analyse the frequency of observation of the HA mutants in the competitive replication assay.

**Supplementary Table S1.** Prevalence of amino acids at indicated positions of the H3 HA of avian IAVs

**Supplementary Table S2.** Originating and submitting laboratories of the sequences from GISAID’s EpiFlu™ Database on which this research is based.

**Supplementary Fig. S1.** Inference of amino acid substitutions separating HAs of H3N2/1968 pandemic IAVs from their avian ancestor.

**Supplementary Fig. S2.** Evolution of glycosylation site at HA positions 63 and 81 of human H3N2 IAVs.

**Supplementary Fig. S3.** Conformational stability and membrane-fusion properties of the HA point mutants of HK and R5.

**Supplementary Fig. S4.** Receptor-binding properties of HK, R5 and their mutants grown in Calu-3 cells and HTBE cultures.

**Supplementary Fig. S5.** Variation of amino acids at 9 positions of H3 HA during evolution in different host species.

**Supplementary Fig. S6.** Amino-acid composition at the nine sites of H3 HA.

## Acknowledgements

We thank Earl Brown for A/Hong Kong/1/1968 (H3N2) and Robert Webster and Richard Webby for avian influenza viruses and pHW2000 and PR8 plasmids. We gratefully acknowledge GISAID Initiative and all authors from the originating and submitting laboratories of the sequences in the GISAID’s EpiFlu Database on which part of this research is based. The list is detailed in the supplementary Table S2.

This work was supported by the European Union’s Seventh Framework Programme for Research, Technological Development, and Demonstration under grant agreement 278433-PREDEMICS, by the Deutsche Forschungsgemeinschaft (DFG, German Research Foundation) - Project number 197785619 – SFB 1021, by an NWO VIDI grant (contract number 91715372) and by NIH/NIAID contract HHSN272201400008C. Johanna West was supported by a fellowship from the Jürgen Manchot Stiftung, Düsseldorf, Germany.

## Author contributions

**Conceptualization:** Hans-Dieter Klenk, Mikhail Matrosovich

**Formal Analysis:** Gianpiero Zamperin, Michele Gastaldelli, Francesco Bonfante, Jochen Wilhelm, Mikhail Matrosovich

**Funding Acquisition**: Johanna West, Sander Herfst, Ron Fouchier, Mikhail Matrosovich

**Investigation:** Johanna West, Juliane Röder, Tatyana Matrosovich, Jana Beicht, Jan Baumann, Nancy Mounogou Kouassi, Jennifer Doedt, Annalisa Salviato, Sergei Kosakovsky Pond, Sander Herfst

**Methodology:** Gianpiero Zamperin, Michele Gastaldelli, Francesco Bonfante, Sergei Kosakovsky Pond, Sander Herfst, Ron Fouchier, Jochen Wilhelm, Mikhail Matrosovich

**Resources:** Nicolai Bovin

**Supervision:** Mikhail Matrosovich

**Visualization:** Johanna West, Sergei Kosakovsky Pond, Sander Herfst, Jochen Wilhelm, Mikhail Matrosovich

**Writing – Original Draft:** Johanna West, Mikhail Matrosovich

**Writing – Review & Editing:** Johanna West, Nicolai Bovin, Francesco Bonfante, Sergei Kosakovsky Pond, Sander Herfst, Ron Fouchier, Jochen Wilhelm, Hans-Dieter Klenk, Mikhail Matrosovich

