## Supplementary Method for "Characterization of changes in the hemagglutinin that accompanied the emergence of H3N2/1968 pandemic influenza viruses"

**Supplementary Method. Construction of a GAMLSS model to analyse the frequency of observation of the HA mutants in the competitive replication assay.**

In order to determine the effect of the inoculum titre on the frequency of each SAP, we employed a generalized linear model for location, scale and shape (GAMLSS model). relating the observed frequencies to the inoculum titre (expressed by the variable “treatment”), the locus of each SAP (expressed by the variable “SAP”) and the interaction of these two variables. The variable “treatment” consisted of the levels “control”, “L”, “M”, and “H”, while “SAP” was described by the levels “L15F”, “I62R”, “N81D”, “K92N”, “G144A” and “S193N” GAMLSS models allow modeling not only the mean, but also location and scale parameters of the dependent variable distribution as linear parametric functions of explanatory variables. The model was constructed under R environment(R Core Team, 2018) and, in order to deal with the overdispersion of the frequency values, it assumed a beta-binomial distribution in which the dispersion parameter σ was estimated as a function of “treatment” (Rigby & Stasinopoulos, 2005).

In GAMLSS models, the beta binomial distribution is depicted as BB(n, µ, σ), where n is the number of observations, µ is the event probability and is comprised in the range 0 < µ < 1 and σ is the dispersion parameter and is > 0. The mean E(Y) and variance Var(Y) are respectively expressed as nµ and $n\mu\left( 1-\mu\right)\left[ 1+\frac{\sigma}{1+\sigma}\left( n-1 \right) \right]$.

The probability density function of the beta binomial distribution as BB(n, µ, σ) is given by

$$p_{Y}\left( y|\mu, \sigma\right)= \frac{\Gamma\left( n+ 1 \right)}{\Gamma\left( y+1 \right)\Gamma\left( n-y+1 \right)}\frac{\Gamma\left( \frac{1}{\sigma} \right)\Gamma\left( 1+\frac{\mu}{\sigma} \right)\Gamma\left( n+\frac{\left( 1-\mu\right)}{\sigma}-y \right)}{\Gamma\left( n+\frac{1}{\sigma} \right)\Gamma\left( \frac{\mu}{\sigma} \right)\Gamma\left( \frac{1-\mu}{\sigma} \right)}$$

Inside the model, the variable “treatment” was coded by the dummy variables “trtL”, “trtM” and “trtH” which assumed values 1 when the “treatment” level was “L”, “M”, and “H” respectively and 0 otherwise. The “control” level was described by assigning the value 0 to the dummy variables. Similarly, the variable “SAP” was expressed by the dummy variables “L15F”, “I62R”, “N81D”, “K92N” and “S193N” which assumed value 1 when the described SAP was L15F, I62R, N81D, K92N and S193N respectively, and 0 otherwise. G144A was described by assigning the value 0 to the dummy variables.

The final model consisted of 2 parts, the one modelling the frequency $\mu$

$$\log\left( \frac{\mu}{1-\mu} \right)={Intercept}_{\mu}+ \hat{\beta}_{\mu\_trtH} trtH+ \hat{\beta}_{\mu\_trtM} trtM+ \hat{\beta}_{\mu\_trtL} trtL+\hat{\beta}_{\mu\_L15F} L15F+ \hat{\beta}_{\mu\_I62R} I62R+ \hat{\beta}_{\mu\_N81D} N81D+ \hat{\beta}_{\mu\_K92N} K92N+\hat{\beta}_{\mu\_S193N} S193N+\hat{\beta}_{\mu\_trtH\_L15F} trtH L15F+\hat{\beta}_{\mu\_trtM\_L15F} trtM L15F+\hat{\beta}_{\mu\_trtL\_L15F} trtL L15F+ \hat{\beta}_{\mu\_trtH\_I62R} trtH I62R+ \hat{\beta}_{\mu\_trtM\_I62R} trtM I62R+ \hat{\beta}_{\mu\_trtL\_I62R} trtL I62R+ \hat{\beta}_{\mu\_trtH\_N81D} trtH N81D+ \hat{\beta}_{\mu\_trtM\_N81D} trtM N81+ \hat{\beta}_{\mu\_trtL\_N81D} trtL N81+ \hat{\beta}_{\mu\_trtH\_K92N} trtH K92N+ \hat{\beta}_{\mu\_trtM\_K92N} trtM K92N+ \hat{\beta}_{\mu\_trtL\_K92N} trtL K92N+ \hat{\beta}_{\mu\_trtH\_S193N} trtH S193N+ \hat{\beta}_{\mu\_trtM\_S193N} trtM S193N+ \hat{\beta}_{\mu\_trtL\_S193N} trtL S193N$$

and the one modelling the dispersion parameter $\sigma$

$$\log\left( \sigma\right)={Intercept}_{\sigma}+ \hat{\beta}_{\sigma\_trtH} trtH+ \hat{\beta}_{\sigma\_trtM} trtM+ \hat{\beta}_{\sigma\_trtL} trtL$$

The model parameter estimates are given in the Table A. The significance of each predictive variable for µ and σ was assessed via likelihood ratio test (Table B).

**Table A. Parameter estimates (Est) and 95% confidence intervals (95% CI) of the GAMLSS model for the analysis of the frequency of observation of the HA mutants in the competitive replication assay.**

| **Frequency modelling** | |  |  |
| --- | --- | --- | --- |
| *Predictors* | *Est* | | *95% CI* |
| $\mathrm{Intercept}_{\mu}$ | -0.56 | | (-0.69, -0.43) |
| $\hat{\beta}_{\mu\_trtH}$ | -0.27 | | (-0.53, -0.01) |
| $\hat{\beta}_{\mu\_trtM}$ | -0.35 | | (-1.23, 0.52) |
| $\hat{\beta}_{\mu\_trtL}$ | -0.99 | | (-2.16, 0.19) |
| $\hat{\beta}_{\mu\_L15F}$ | -2.07 | | (-2.37, -1.78) |
| $\hat{\beta}_{\mu\_I62R}$ | -1.18 | | (-1.41, -0.96) |
| $\hat{\beta}_{\mu\_N81D}$ | -0.99 | | (1.21, -0.78) |
| $\hat{\beta}_{\mu\_K92N}$ | -1.40 | | (-1.63, -1.16) |
| $\hat{\beta}_{\mu\_S193N}$ | -1.60 | | (-1.85, -1.35) |
| $\hat{\beta}_{\mu\_trtH\_L15F}$ | 0.91 | | (0.43, 1.40) |
| $\hat{\beta}_{\mu\_trtM\_L15F}$ | 1.62 | | (0.37, 2.88) |
| $\hat{\beta}_{\mu\_trtL\_L15F}$ | 2.36 | | (0.86, 3.86) |
| $\hat{\beta}_{\mu\_trtH\_I62R}$ | 0.11 | | (-0.33, 0.54) |
| $\hat{\beta}_{\mu\_trtM\_I62R}$ | -0.47 | | (-1.90, 0.95) |
| $\hat{\beta}_{\mu\_trtL\_I62R}$ | -0.50 | | (-2.41, 1.42) |
| $\hat{\beta}_{\mu\_trtH\_N81D}$ | -0.18 | | (-0.61, 0.26) |
| $\hat{\beta}_{\mu\_trtM\_N81D}$ | -0.21 | | (-1.53, 1.12) |
| $\hat{\beta}_{\mu\_trtL\_N81D}$ | -0.81 | | (2.72, 1.11) |
| $\hat{\beta}_{\mu\_trtH\_K92N}$ | 0.18 | | (-0.28, 0.63) |
| $\hat{\beta}_{\mu\_trtM\_K92N}$ | -0.01 | | (-1.38, 1.37) |
| $\hat{\beta}_{\mu\_trtL\_K92N}$ | -0.06 | | (-1.86, 1.73) |
| $\hat{\beta}_{\mu\_trtH\_S193N}$ | 0.25 | | (-0.22, 0.72) |
| $\hat{\beta}_{\mu\_trtM\_S193N}$ | -0.09 | | (-1.48, 1.29) |
| $\hat{\beta}_{\mu\_trtL\_S193N}$ | 1.31 | | (-0.25, 2.88) |
| **Sigma modelling** |  | |  |
| *Predictors* | *Est* | | *95% CI* |
| $\mathrm{Intercept}_{\sigma}$ | -7.10 | | (-7.19, -7.02) |
| $\hat{\beta}_{\sigma\_trtH}$ | 2.11 | | (1.94, 2.27) |
| $\hat{\beta}_{\sigma\_trtM}$ | 5.89 | | (5.32, 6.45) |
| $\hat{\beta}_{\sigma\_trtL}$ | 6.75 | | (5.99, 7.51) |

**Table B.** **Analysis of deviance table (type II likelihood ratio tests) of the GAMLSS model for the analysis of the frequency of observation of the HA mutants in the competitive replication assay**. The model relates the frequency to the variables “treatment”, “SAP” and their interaction, and the dispersion parameter sigma to “treatment” alone.

| **Frequency modelling** |  |  |  |  |
| --- | --- | --- | --- | --- |
| *Predictors* | *npar* | *AIC* | *Χ ^2^* | *p* |
| <none> |  | 2617.1 |  |  |
| treatment:SAP | 15 | 2650.2 | 63.17 | 7.14x10^-8^ |
| **Sigma modelling** |  |  |  |  |
| *Predictors* | *npar* | *AIC* | *Χ ^2^* | *p* |
| Treatment | 3 | 2786.8 | 175.79 | 7.15x10^-38^ |

Legend: *npar =* number of parameters associated to the relative predictor; *AIC =* Akaike’s information criterion value observed upon dropping of the relative predictor; *Χ ^2^ =* likelihood ratio test statistic; p = likelihood ratio test *p* value.
