## Supplementary Table S1 for "Characterization of changes in the hemagglutinin that accompanied the emergence of H3N2/1968 pandemic influenza viruses"

**Supplementary Table S1. Prevalence of amino acids at indicated positions of the H3 HA of avian IAVs[a]**

| Position[b] | A | C | D | E | F | G | H | I | K | L | M | N | P | Q | R | S | T | V | W | Y |
|---|---|---|---|---|---|---|---|---|---|---|---|---|---|---|---|---|---|---|---|---|
| **-2(15)** | . | . | . | . | 946[c] | . | . | 3 | . | 443[c] | . | . | . | . | . | 95 | 1 | 1 | . | 5 |
| **62(78)** | . | . | . | . | . | 3 | . | . | 7 | . | . | . | . | . | 1484 | . | . | . | . | . |
| **63(79)** | 1 | . | 1468 | 4 | . | 17 | . | . | . | . | . | 1 | . | . | . | 1 | . | . | . | . |
| *64(80)* | . | *1494* | . | . | . | . | . | . | . | . | . | . | . | . | . | . | . | . | . | . |
| *65(81)* | *14* | . | . | . | . | . | . | . | *1* | . | . | . | . | . | *4* | *1* | *1474* | . | . | . |
| **81(97)** | . | . | 1473 | 3 | . | 13 | 2 | . | . | . | . | 1 | . | . | . | . | . | . | . | 2 |
| *82(98)* | *26* | . | . | *1446* | . | *1* | . | . | *5* | . | *2* | . | . | . | *1* | . | . | *13* | . | . |
| *83(99)* | *1* | . | . | . | . | . | . | *1* | . | . | *3* | *5* | . | . | . | . | *1484* | . | . | . |
| **92(108)** | . | . | . | 1 | 1 | 4 | 2 | 1 | . | . | . | 1002 | . | . | . | 483 | . | . | . | . |
| **144(160)** | 1319 | . | 10 | . | . | . | . | . | . | . | . | 1 | . | . | . | 1 | 134 | 29 | . | . |
| **193(209)** | . | . | 116 | . | . | 1 | . | . | 6 | . | . | 1070 | . | . | 2 | 297 | 2 | . | . | . |
| **226(242)** | . | . | . | . | . | . | . | . | . | . | . | . | . | 1494 | . | . | . | . | . | . |
| **228(244)** | . | . | . | . | . | 1494 | . | . | . | . | . | . | . | . | . | . | . | . | . | . |

[a] Full-length HA sequences of H3 subtype avian IAVs were downloaded from the GISAID EpiFlu database and processed as described in the legend to supplementary Fig. S1. The final dataset contained 1494 unique full-length sequences. The table shows numbers of sequences with indicated amino acid at each position calculated using Bio-Edit version 7.1.11. Dot is used instead of zero for clarity.

[b] The first number shows amino acid position in the mature HA sequence. The number in parentheses corresponds to the position of the amino acid within the complete coding sequence of the HA precursor. The numbers of 9 HA positions separating pandemic viruses from the avian precursor are shown by bold characters; positions 64-65 and 82-83 (*italics*) represent 2$^{nd}$ and 3$^{rd}$ position of the glycosylation sites $N_{63}$-$C_{64}$-$T_{65}$ and $N_{81}$-$E_{82}$-$T_{83}$ in the human HAs.

[c] *Cyan* and *yellow* colours depict amino acids found in the H3N2/1968 pandemic IAVs and in their inferred avian precursor, respectively. In particular, human-type amino acids $I_{62}$, $K_{92}$, $G_{144}$, $L_{226}$ and $S_{228}$ were not found in any of the avian HA sequences analysed. Only two avian H3N8 viruses, A/blue-winged teal/Missouri/15OS4859/2015 and A/duck/Ukraine/1/1963, contained $N_{63}$ and $N_{81}$, respectively, which encoded N-glycosylation sites $N_{63}$-$C_{64}$-$T_{65}$ and $N_{81}$-$E_{82}$-$T_{83}$.
