## Supplementary Figures S1-S6 for "Characterization of changes in the hemagglutinin that accompanied the emergence of H3N2/1968 pandemic influenza viruses"

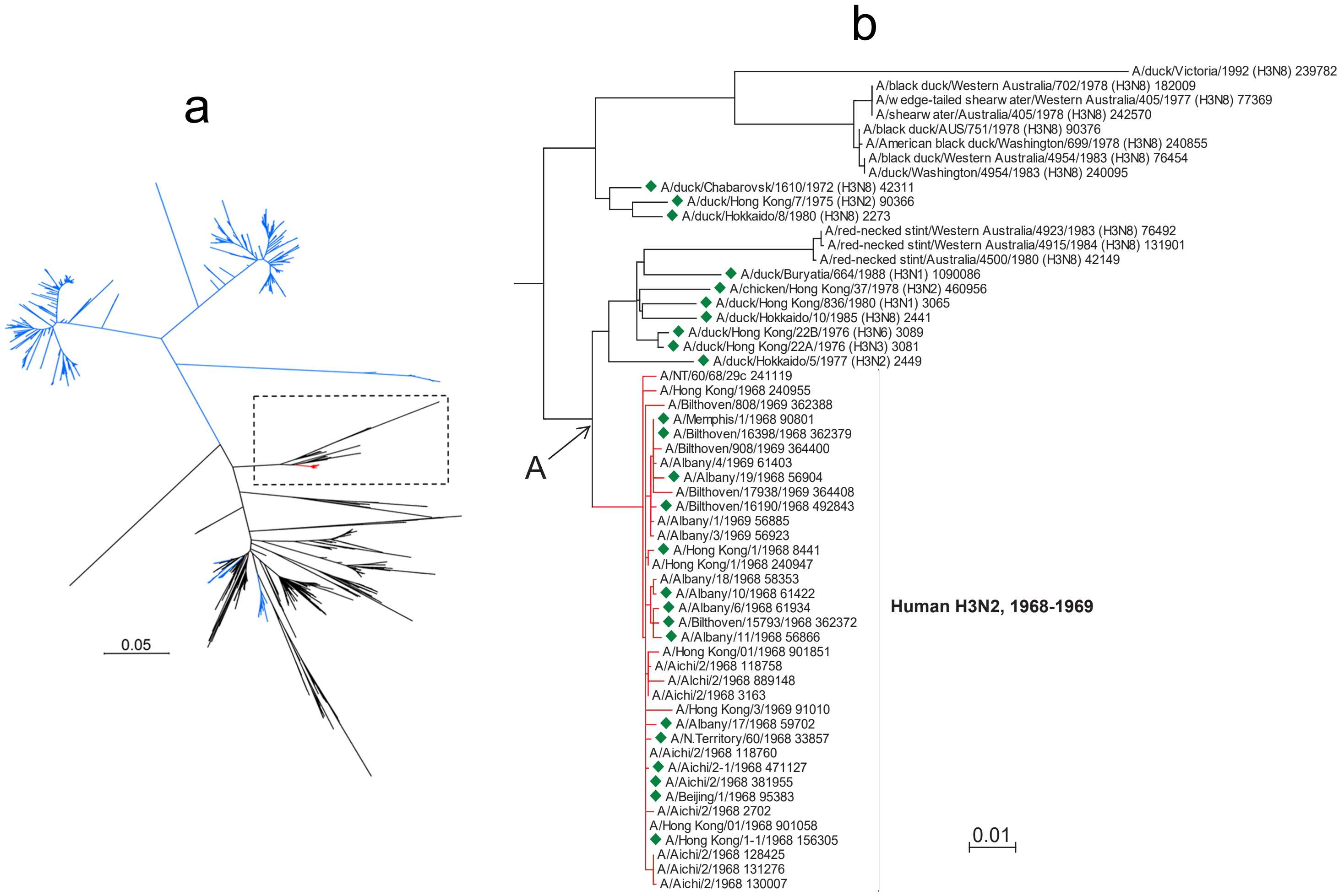


**Supplementary Fig. S1. Inference of amino acid substitutions separating HAs of H3N2/1968 pandemic IAVs from their avian ancestor.**

(**a**) Phylogenetic relationships between H3N2/1968 and avian IAVs. All full-length HA sequences of avian H3 IAVs and of pandemic IAVs isolated in 1968-1969 were downloaded from the GISAID EpiFlu database accessed on March 11, 2020. The sequences were aligned using MAFFT program implemented in the Unipro UGENE package (Okonechnikov et al., 2012), version 1.32. Sequences containing gaps and ambiguities, non-unique sequences and sequences of swine-origin avian and laboratory-derived IAVs were removed manually using Bio-Edit version 7.1.11 (Hall, 1999). The final dataset contained sequences of 1494 avian and 36 human HAs. The evolutionary history was inferred using MEGA7 (Kumar et al., 2016) with the maximum likelihood method based on the Kimura 2-parameter model. The tree is drawn to scale, with branch lengths measured in the number of nucleotide substitutions per site. Colours depict avian IAVs from North America (*blue*), Eurasia and Oceania (*black*), and H3N2/1968 pandemic viruses (*red*).

(**b**) Detailed view of the branch, which includes pandemic and the closest avian IAVs marked by the dashed box in the panel a. Subtypes of avian IAVs and accession numbers of all sequences are shown next to the strain names. HA amino acid sequence of the common avian-human ancestor (node A) was inferred using the Ancestors program of MEGA7. The ancestral sequence together with the sequences of representative avian and human IAVs depicted by *green diamonds* are shown in the panel c.

(**c**) see next page.

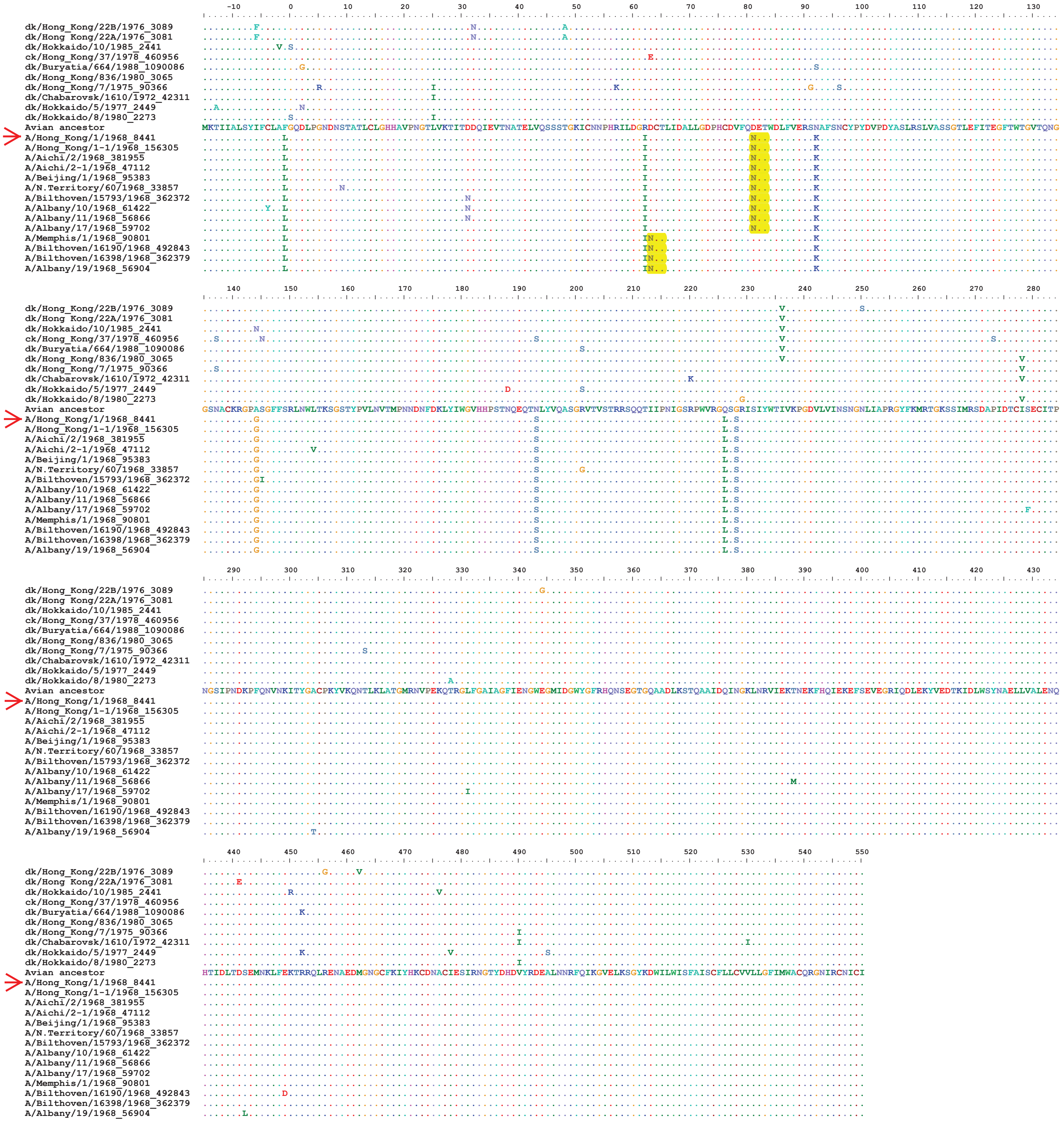


**Supplementary Fig. S1. Inference of amino acid substitutions separating HAs of H3N2/1968 pandemic IAVs from their avian ancestor.**

**(a,b)** see previous page

**(c)** HA amino acid sequences of H3N2 pandemic IAVs viruses isolated in 1968, the closest avian IAVs and the inferred common avian-human ancestor. Numbering starts from the N-terminus of mature HA protein, the signal peptide is numbered from -15 to 0. Dots depict identity with the sequence of the avian ancestor. Glycosylation sites at HA positions 63-65 and 81-83 are highlighted by *yellow*. *Arrow* depicts virus strain A/Hong Kong/1/1968 (H3N2) used to make recombinant viruses in this study. The figure was generated using Bio-Edit.

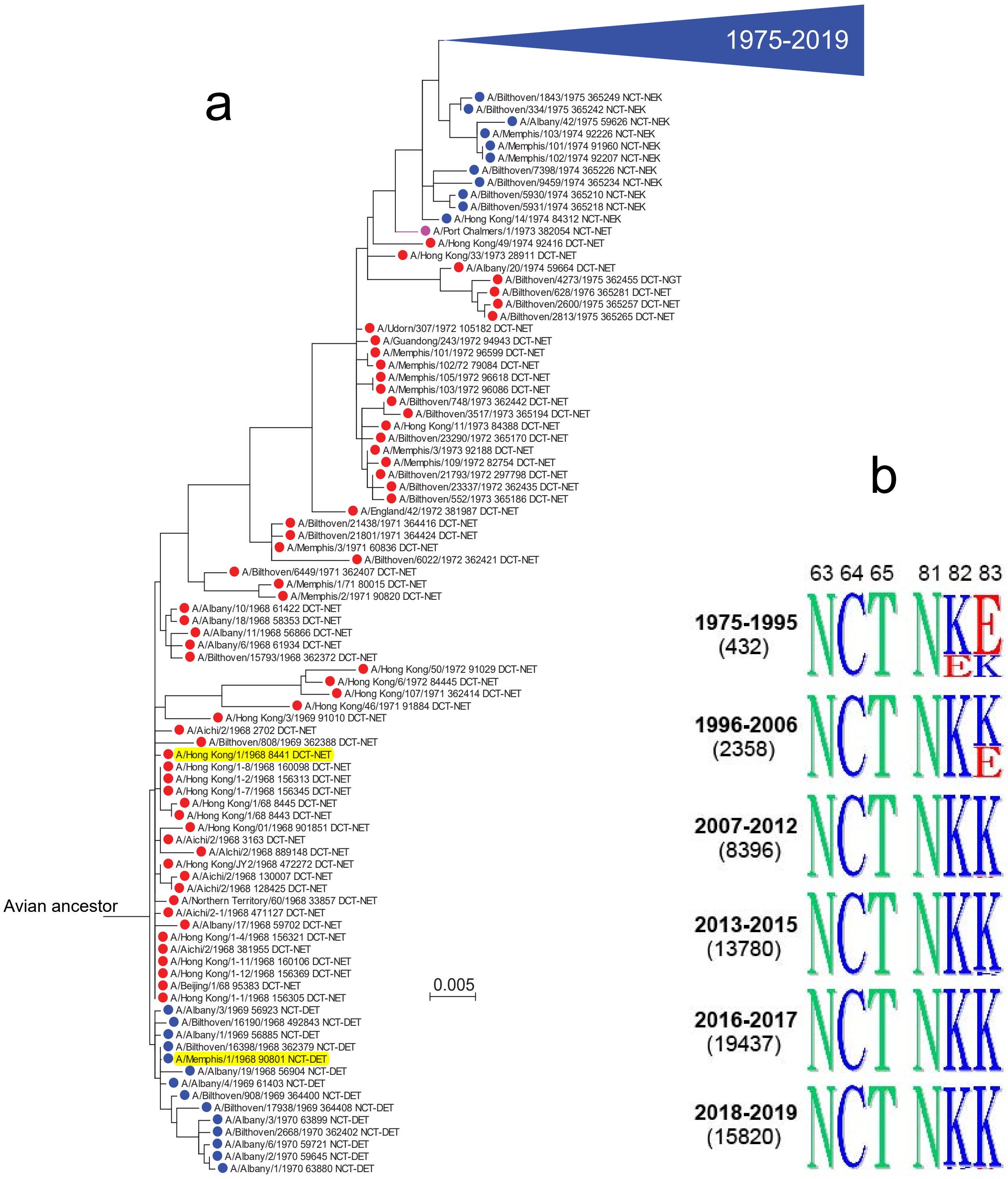


**Supplementary Fig. S2. Evolution of glycosylation sites in HA positions 63 and 81 of human H3N2 IAVs.** Full-length HA sequences were downloaded and processed as described in the legend to supplementary Fig. S1.

(**a**) Phylogenetic relationships between the HA of IAVs isolated from 1968 to 1995 were inferred using MEGA7 with the maximum likelihood method. GISAID accession numbers and amino acids in positions 63-65 and 81-83 are shown next to the strain names. *Coloured circles* depict presence of glycosylation sites 63 (*blue*) and 81 (*red*). One virus strain (A/Port Chalmers/1/1973) contained glycosylation sites at both positions (*magenta*). The branch containing sequences of the viruses isolated after 1975 is collapsed for clarity (*blue triangle*). Prototype strains used to define two glycosylation lineages of pandemic viruses are highlighted by *yellow*.

(**b**) Protein logos show frequencies of amino acids at positions 63-65 and 81-83 of the HA. Years of virus isolation and numbers of analysed sequences (in parentheses) are shown on the left. The figure was generated using Phylo-mLogo software (Shih et al., 2007).

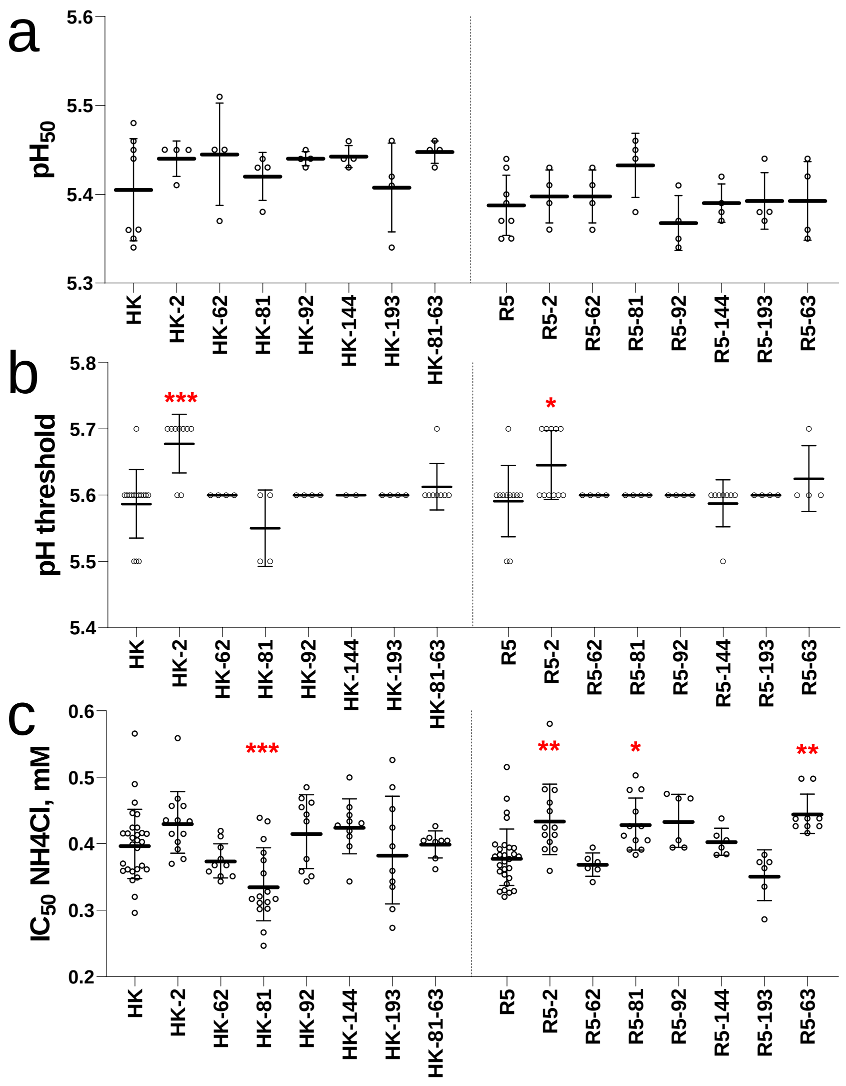


**Supplementary Fig. S3. Conformational stability and membrane-fusion properties of the HA point mutants of HK and R5.**

(**a**) pH of acid-induced conformational transition. Solid-phase adsorbed viruses were incubated in acidic buffers for 15 min at 37oC, returned to PBS and treated with proteinase K. Viral binding of peroxidase-labeled fetuin was assayed, and pH values that corresponded to 50% reduction of HA binding activity ($pH_{50}$) were determined from binding-versus-pH curves.

(**b**) pH threshold of polykaryon formation. MDCK cells were infected at m.o.i. 1, cultured overnight, treated with trypsin and exposed to different pH for 10 min at 37°C. After returning to neutral medium and incubation for 3 h the cells were fixed, stained with Giemsa stain and analysed under the microscope. The data show highest pH values at which formation of polykaryons was detected.

(**c**) Inhibition of viral infection by ammonium chloride. MDCK cells were infected in the presence of various concentrations of $NH_4Cl$, incubated overnight, fixed, and immune-stained for NP. Concentrations of $NH_4Cl$ that reduced numbers of infected cells by 50% ($IC_{50}$) were determined from dose-response curves. Figures show data points, geometric mean and SDs from 2 to 8 experiments performed on different days with 2 to 6 replicates each. Data in panela a and c were analysed using general linear mixed models in R 3.6.0. Concentration was log-transformed before analysis. Day was included as a random intercept.

Figures show data adjusted for day. In panel b, due to the low resolution of the assay (with only 3 distinct pH values), pH was analysed as an ordinal variable using an ordered logistic regression model. Multiple tests of contrasts within these models were done using simultaneous tests for general linear hypotheses, and P values were adjusted using the single step method. In all panels, vertical dotted line separates point mutants of HK and point mutants of R5. *Red* asterisks depict point mutants that were significantly different from their parental viruses, either HK or R5. No significant difference was observed between HK and R5 in these assays.

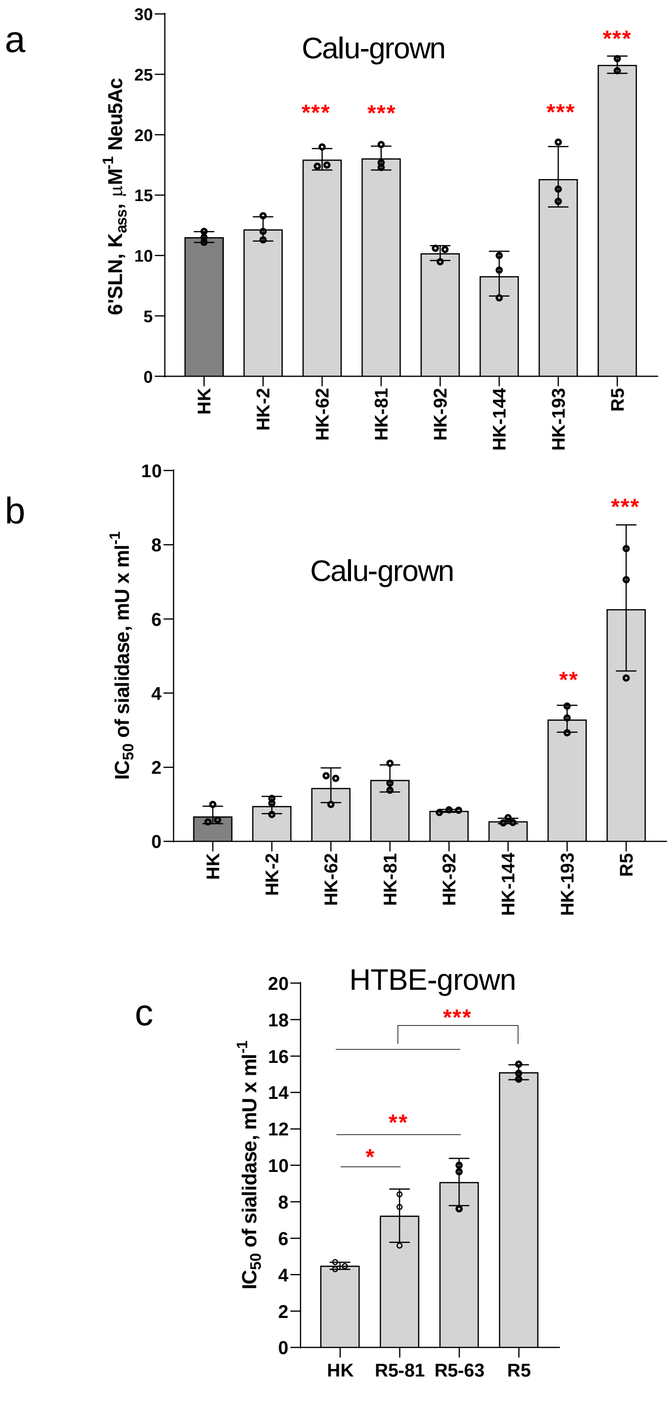


**Supplementary Fig. S4. Receptor-binding properties of HK, R5 and their mutants grown in Calu-3 cells and HTBE cultures.**

(**a**) Association constants of viral complexes with 6'SLN (20 kDa).

(**b,c**) Virus avidity for receptors on MDCK cells expressed as concentrations of the *Vibrio cholerae* sialidase that reduced numbers of infected cells by 50% ($IC_{50}$). The higher $IC_{50}$, the higher binding avidity. All panels show replicates, mean values (bars) and SDs. Asterisks in panels a and b show P values for the differences with HK (*dark grey bars*) determined using Dunnett's multiple comparisons test. Asterisks in panel c depict differences between individual viruses determined using Tukey's multiple comparisons test. *, P<0.05; **, P<0.01; ***, P< 0.001.

**Supplementary Fig. S5** (see next page, scalable vector graphics)**. Variation of amino acids at 9 positions of H3 HA during evolution in different host species** . The figure shows maximum likelihood tree for the nucleotide sequences of H3 HA inferred using IQ-TREE 2 (Kalyaanamoorthy et al., 2017; Minh et al., 2020) and plotted using Mega 7 (Kumar et al., 2016). The tree is based on the following non-redundant full-length H3 HA sequences of IAVs from the GISAID EpiFlu database. Representative human IAVs isolated from 1968 to 2020 (106 sequences, 2 per year), representative swine IAVs (456 sequences selected from the total 2663 sequences using JalView (Waterhouse et al., 2009)), all other mammalian IAVs, mainly equine and canine (402 sequences), all avian IAVs (1494 sequences), avian, swine and human IAVs sporadically isolated from a different host species (31 sequences). *Taxon labels* (for example, A/equine/Miami/1/1963-A/H3N8-129722-H-R-NCT-YEN-S-A-K-QSG) include virus name, subtype, GISAID EpiFlu accession number and amino acids in HA positions -2, 62, 63-65, 81-83, 92, 144, 193, 226-228. Colours depict the following stable host-specific lineages. *Black*, avian; *red*, human; *green*, swine; *blue*, equine; *cyan*, equine/Jilin/1/1989; *orange*, canine H3N8; *magent*a, canine H3N2. The earliest virus isolates from mammalian lineages are highlighted with *yellow*. *Green dots* depict 10 sporadic avian-like mammalian isolates.

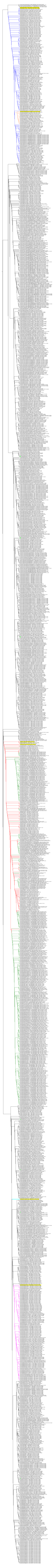

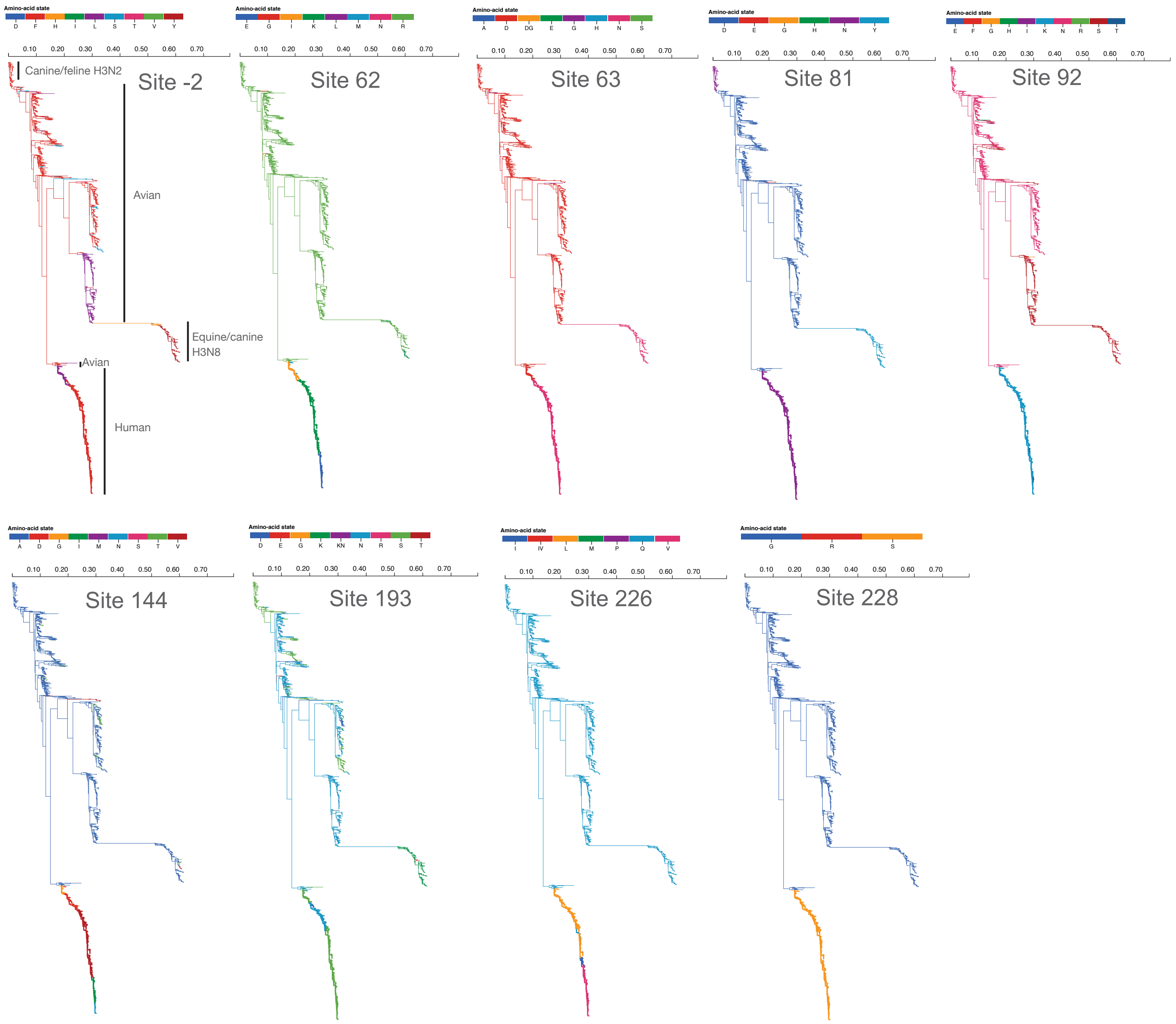


**Supplementary Fig. S6. Amino acid composition at the nine sites of H3 HA** (scalable vector graphics). The tree is based on HA sequences used for selection pressure analyses and includes 1492 sequences of avian IAVs, 406 sequences of equine, canine, feline and seal IAVs and 803 sequences of human IAVs isolated in the years from 1968 to 1999. Unobserved ancestral codons were inferred using the SLAC method. Host-specific clades are depicted in the tree display for the site -2.
